# CDR2 is a dynein adaptor recruited by kinectin to regulate ER sheet organization

**DOI:** 10.1101/2024.11.06.622207

**Authors:** Vanessa Teixeira, Kashish Singh, José B. Gama, Ricardo Celestino, Ana Xavier Carvalho, Paulo Pereira, Carla M.C. Abreu, Tiago J. Dantas, Andrew P. Carter, Reto Gassmann

## Abstract

The endoplasmic reticulum (ER) relies on the microtubule cytoskeleton for distribution and re-modelling of its extended membrane network, but how microtubule-based motors contribute to ER organization remains unclear. Using biochemical and cell-based assays, we identify cerebellar degeneration-related protein 2 (CDR2) and its paralog CDR2-like (CDR2L), onconeural antigens with poorly understood functions, as ER adaptors for cytoplasmic dynein-1 (dynein). We demonstrate that CDR2 is recruited by the integral ER membrane protein kinectin (KTN1) and that double knockout of CDR2 and CDR2L enhances KTN1-dependent ER sheet stacking, reversal of which by exogenous CDR2 requires its dynein-binding CC1 box motif. Exogenous CDR2 expression additionally promotes CC1 box-dependent clustering of ER sheets near centrosomes. CDR2 competes with the eEF1Bβ subunit of translation elongation factor 1 for binding to KTN1, and eEF1Bβ knockdown increases endogenous CDR2 levels on ER sheets, inducing their centrosome-proximal clustering. Our study describes a novel molecular pathway that implicates dynein in ER sheet organization and may be involved in the pathogenesis of paraneoplastic cerebellar degeneration.

## INTRODUCTION

Tight regulation of organelle positioning is a prerequisite for cell health (Barlan and Gelfand, 2017). Organelles are distributed in part through transport along microtubules by the predominantly plus end-directed kinesins and minus end-directed cytoplasmic dynein-1 (dynein). Elucidating how these motors are recruited and activated on membranes to drive bi-directional transport requires the identification and characterization of cargo-specific adaptor proteins (Cross and Dodding, 2019), whose inventory remains incomplete.

The endoplasmic reticulum (ER) is a highly dynamic organelle, yet the recruitment mechanisms and functions of ER-associated microtubule motors are poorly understood (Perkins and Allan, 2021). The ER extends from the nuclear envelope as an interconnected network of sheets and tubules. Sheets are flat cisternal structures that can be arranged into stacks and are enriched in the perinuclear region, whereas ER tubules form a reticular network that is present in both the perinuclear and peripheral regions (Goyal and Blackstone, 2013; Lin *et al*., 2012; Park and Blackstone, 2010; Zhang and Hu, 2016). Sheets typically contain ribosomes (rough ER) and are the site of co-translational translocation of integral membrane and secretory proteins into the ER lumen. Tubules tend to be devoid of ribosomes (smooth ER) and are involved in lipid synthesis and delivery, establishing contact with other organelles, calcium homeostasis, and detoxification. In line with functional specialization of ER subdomains, the proportion of sheets to tubules, as well as their spatial arrangement, can differ significantly depending on cell type and growth conditions. Abnormalities in ER organization are linked to various diseases, including neurodegenerative disorders (Perkins and Allan, 2021; Goyal and Blackstone, 2013; Westrate *et al*., 2015).

Kinesin-1 and dynein associate with microsomes isolated from brain (Yu *et al*., 1992), and both motors have been implicated in ER dynamics, primarily the movement of tubules (Allan and Vale, 1991; Allan, 1995; FitzHarris *et al*., 2007; Lane and Allan, 1999; Mukherjee *et al*., 2020; Niclas *et al*., 1996; Steffen *et al*., 1997; Wang *et al*., 2013; Wedlich-Söldner *et al*., 2002; Woźniak *et al*., 2009). Kinectin (KTN1), an integral membrane protein that is enriched on ER sheets (Shibata *et al*., 2010), was the first membrane receptor for kinesin-1 to be identified (Fütterer *et al*., 1995; Kumar *et al*., 1995; Toyoshima *et al*., 1992). KTN1 binds the C-terminus of kinesin heavy chain KIF5 via its extended cytosolic coiled-coil domain (Ong *et al*., 2000). The KTN1 paralog RRBP1 (p180), a receptor for ribosomes on ER membranes (Koppers *et al*., 2024; Savitz and Meyer, 1990; Ueno *et al*., 2012; Wanker *et al*., 1995), binds KIF5 in an analogous manner (Diefenbach *et al*., 2004). The KTN1–kinesin-1 interaction is proposed to promote ER transport to the cell periphery to support focal adhesion growth and maturation (Guadagno *et al*., 2020; Ng *et al*., 2016; Santama *et al*., 2004; Zhang *et al*., 2010).

Dynein adaptors on ER membranes have yet to be identified. The mega-dalton dynein complex is built around a dimer of the heavy chain (DHC) that contains the motor domain at its C-terminus and interacts with intermediate and light intermediate chains (DIC and DLIC, respectively) via its N-terminal region (Canty *et al*., 2021; Carter *et al*., 2016). In recent years, several so-called activating adaptors have been identified that form a tripartite complex with the dynein N-terminus and the obligatory dynein co-factor dynactin, also a mega-dalton complex (McKenney *et al*., 2014; Schlager *et al*., 2014; Urnavicius *et al*., 2015, 2018). Activating adaptors are characterized by a dimeric N-terminal coiled-coil that stabilizes the dynein–dynactin complex through interactions that are similar across adaptor families (Chabaan and Carter, 2022; Urnavicius *et al*., 2015, 2018; Singh *et al*., 2024), while the more divergent C-termini connect the processive transport machine to specific cargo, for example via binding to membrane-associated proteins (Olenick and Holzbaur, 2019; Reck-Peterson *et al*., 2018). In all activating adaptors identified to date, the N-terminus binds a conserved C-terminal helix in DLIC, and this interaction is important for dynein motility *in vitro* and dynein function in cells (Celestino *et al*., 2019; Lee *et al*., 2018; Schroeder and Vale, 2016). Some adaptors use a short coiled-coil segment referred to as the CC1 box to bind the DLIC helix (Gama *et al*., 2017; Lee *et al*., 2020). Additional contact with dynein occurs through the heavy chain binding site 1 (HBS1) of the adaptor, located approximately 30 residues downstream of the CC1 box (Chabaan and Carter, 2022; Sacristan *et al*., 2018).

Here, we identify cerebellar degeneration-related protein 2 (CDR2) and its paralog CDR2-like (CDR2L) as CC1 box-and HBS1-containing proteins that bind dynein–dynactin, and we demonstrate that purified CDR2L activates dynein motility *in vitro*. CDR2 and CDR2L are associated with paraneoplastic cerebellar degeneration (PCD), a rare immune-mediated disorder triggered by gynaecological cancers (Abbatemarco *et al*., 2024). In patients with PCD, tumor-induced autoimmunity against neuronal antigens, including CDR2/CDR2L, causes degeneration of Purkinje cells in the cerebellum, but the pathogenetic mechanism and the physiological roles of CDR2 and CDR2L remain unclear (Greenlee and Brashear, 2023). CDR2L associates with ribosomes (Herdlevaer *et al*., 2020; Hida *et al*., 1994; Rodriguez *et al*., 1988), and CDR2 is proposed to be involved in transcriptional regulation (O’Donovan *et al*., 2010; Okano *et al*., 1999; Sakai *et al*., 2001, 2002; Takanaga *et al*., 1998).

We demonstrate that CDR2 and CDR2L localize to ER sheets and describe the underlying molecular interactions. CDR2 is recruited by KTN1 and regulates ER sheet organization via its interaction with KTN1 and dynein. CDR2 competes with eEF1Bβ, a subunit of the translation elongation factor 1 complex (Negrutskii *et al*., 2023) (also known as EF1-δ or EF1D) for KTN1 binding, and we provide evidence that altering the relative abundance of CDR2 and eEF1Bβ on ER sheets impacts their distribution. Our findings, which have potential relevance for the pathogenesis of PCD, establish CDR2 and CDR2L as dynein adaptors for the ER that contribute to ER organization.

## RESULTS AND DISCUSSION

### CDR2 and CDR2L are novel adaptors for cytoplasmic dynein-1

The first coiled-coil of human CDR2 and its paralog CDR2L contain a N-terminal CC1 box followed by an HBS1 (Fig. 1A; Fig. S1A, B), suggesting they might be novel dynein adaptors. To test this, we first determined whether the CC1 box binds DLIC using purified recombinant proteins. Size exclusion chromatography (SEC) demonstrated that CDR2(1–146) forms a complex with GST::DLIC1(388–523), but not when the CC1 box is deleted (Δ23–39) (Fig. 1B) or when DLIC1 residues known to be critical for CC1 box binding are mutated (F447A/F448A) (Celestino *et al*., 2019) (Fig. S1C). GST pull-down experiments likewise showed that CDR2L binds GST::DLIC1(388–523) in a CC1 box-dependent manner (Fig. 1C). We next assessed the ability of purified recombinant CDR2 and CDR2L to form complexes with dynein and dynactin from porcine brain lysate. When affinity-isolated from lysate through their C-terminal Strep-tag II, CDR2(1–146), CDR2L(1–159) and CDR2L(1–290) co-isolated dynein–dynactin (Fig. 1D). CDR2(1–146) performed as robustly in this assay as the previously characterized activating adaptor fragment JIP3(1–185) (Singh *et al*., 2024), while CDR2L fragments were less efficient. The CC1 box in CDR2(1–146) and CDR2L(1–290) was essential for complex formation, and introducing point mutations into the HBS1 motif (HBS1_6A) of CDR2(1–146) reduced complex formation. Finally, we assessed whether CDR2 and CDR2L support processive movement of tetramethylrhodamine (TMR)-labeled dynein–dynactin in a motility assay, using Lis1 to facilitate dynein activation (Baumbach *et al*., 2017) (Fig. 1E). This revealed that CDR2L fragments 1–159 and 1–290 can activate dynein motility in a CC1 box-dependent manner, although they were significantly less potent than JIP3(1–185). Unexpectedly, we failed to detect dynein activation with CDR2(1–146), the reason for which remains unclear. Longer CDR2 fragments also failed to activate, although we note that dynein adaptors typically adopt autoinhibited conformations (d’Amico *et al*., 2022; Hoogenraad *et al*., 2003; Singh *et al*., 2024). Taken together, the results from binding assays and *in vitro* re-constitution of dynein motility suggest that human CDR2 and CDR2L are activating adaptors for dynein. Furthermore, the results show that the CC1 box is essential for formation of dynein–dynactin–CDR2/CDR2L complexes. Consistent with this, the *D. melanogaster* CDR2 homolog Centrocortin was recently shown to require the CC1 box for centrosome-directed transport of its mRNA (Zein-Sabatto *et al*., 2024, *Preprint*).

**Figure 1:**
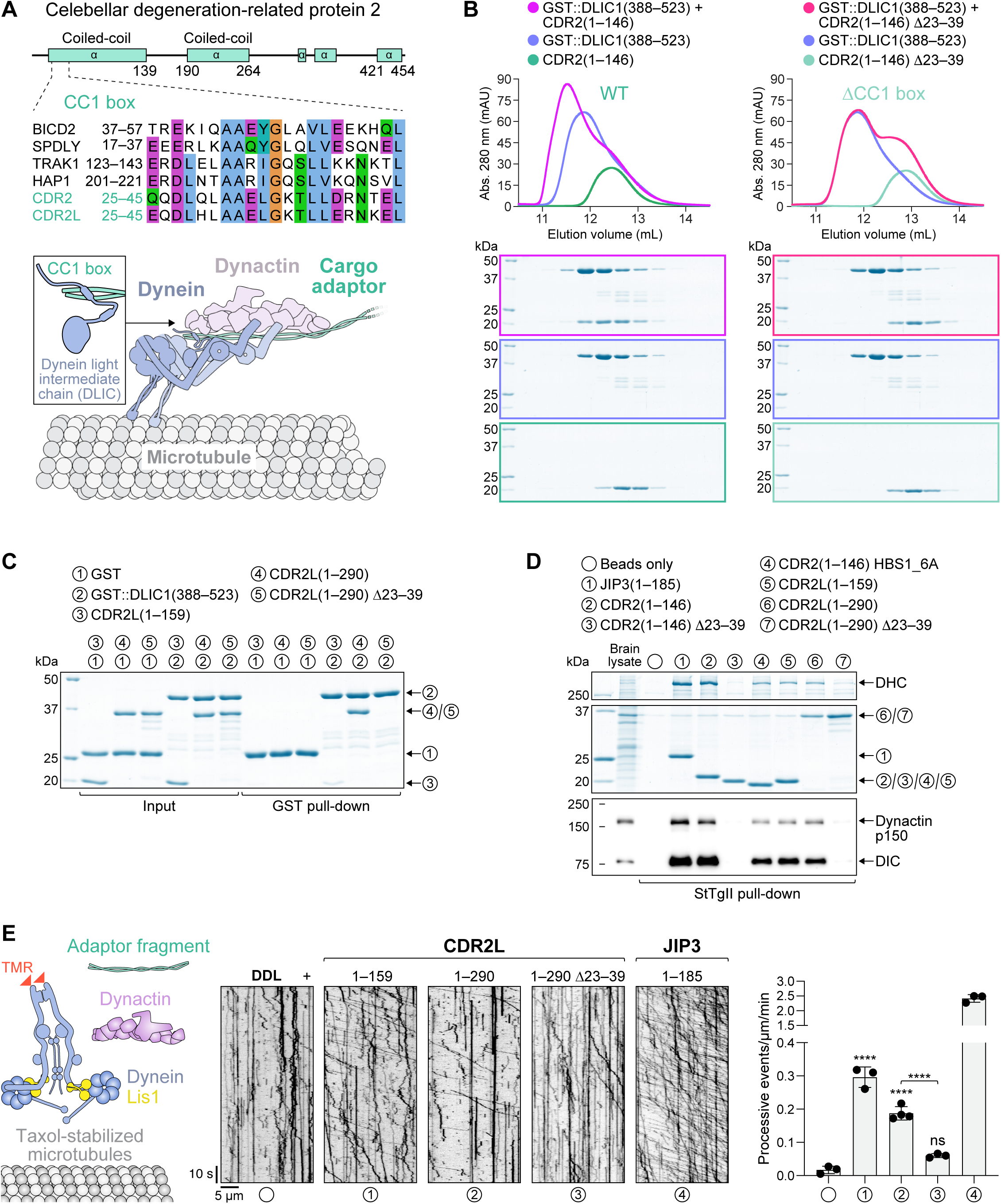
CDR2 and CDR2L are novel adaptors for cytoplasmic dynein-1. **(A)** Schematic of the human CDR2 protein and sequence alignment of its N-terminal CC1 box (motif AAXXG) with that of other human dynein adaptors (see also *Fig. S1B*). The CC1 box binds DLIC, as illustrated in the cartoon below the alignment. **(B)** Elution profiles and BlueSafe-stained SDS-PAGE gels of purified recombinant human CDR2 and DLIC1 fragments after SEC. The elution profile and gel for DLIC1 are shown on both left and right to facilitate comparison between wild-type (WT) CDR2 and the ΔCC1 box mutant. Molecular weight is indicated in kilodaltons (kDa). **(C)** BlueSafe-stained SDS-PAGE gels of purified recombinant proteins prior to addition of glutathione agarose resin (Input) and after elution from the resin (GST pull-down), showing that CDR2L binds to DLIC1. **(D)** BlueSafe-stained SDS-PAGE gel and immunoblot after pull-down of purified recombinant proteins, C-terminally tagged with StTgII, from porcine brain lysate. In the HBS1_6A construct, 6 residues in CDR2’s predicted dynein heavy chain-binding site (HBS1) are mutated to alanine, as shown in *Fig. S1B*. **(E)** *In vitro* motility assays with TMR-labeled dynein, dynactin, Lis1 and adaptor fragments. Representative kymographs and the number of processive events per micrometer of microtubule per minute (mean ± SD of 3-4 technical replicates) are shown. The total number of events analyzed were 21 (DDL), 344 (CDR2L^1-159^), 278 (CDR2L^1-290^), 69 (CDR2L^1-290^ ΔCC1) and 304 (JIP3^1-185^). Statistical significance was determined using ordinary one-way ANOVA followed by Tukey’s multiple comparisons test. \*\*\*\**P* < 0.0001; *ns* = not significant, *P* > 0.05.

### CDR2 and CDR2L interact and co-localize with the integral ER membrane protein KTN1

To identify potential dynein cargo of CDR2 and CDR2L, we performed immunoprecipitations from HeLa cells stably expressing transgenic GFP::3xFLAG-tagged CDR2 or CDR2L in a double knockout background (CDR2/L double KO) (Fig. S2A, B). Quantitative mass spectrometry revealed that the ER sheet component KTN1 was the most enriched protein in anti-FLAG immunoprecipitates from both transgenic cell lines when compared to control immunoprecipitates from parental CDR2/L double KO cells (Fig. 2A). Immunoprecipitates from GFP::3xFLAG::CDR2L cells were additionally enriched for the KTN1 paralog p180. Immunofluorescence revealed striking co-localization of GFP::3xFLAG::CDR2 with KTN1 (Fig. 2B). GFP::3xFLAG::CDR2L also co-localized with KTN1 and in addition exhibited diffuse cytoplasmic localization (Fig. S2C). To test whether the endogenous proteins localize to ER sheets, we co-stained HeLa cells with antibodies against CDR2 and CDR2L and the ER sheet component CLIMP63 (Shibata *et al*., 2010). Endogenous CDR2 was reproducibly detectable on ER sheets by immunofluorescence (Fig. 2C). Signal intensity varied between cells and was generally close to the background signal observed in CDR2 KO cells, which we used as a control for antibody specificity. This suggests CDR2 is expressed at relatively low levels in HeLa cells. Although CDR2L was detectable by immunoblot (Fig. S2A), we were unable to detect specific immunofluorescence signal with multiple antibodies, including our own, against CDR2L. Overall, our data suggests that the dynein adaptors CDR2 and CDR2L interact with ER sheet components and localize to ER sheets.

**Figure 2:**
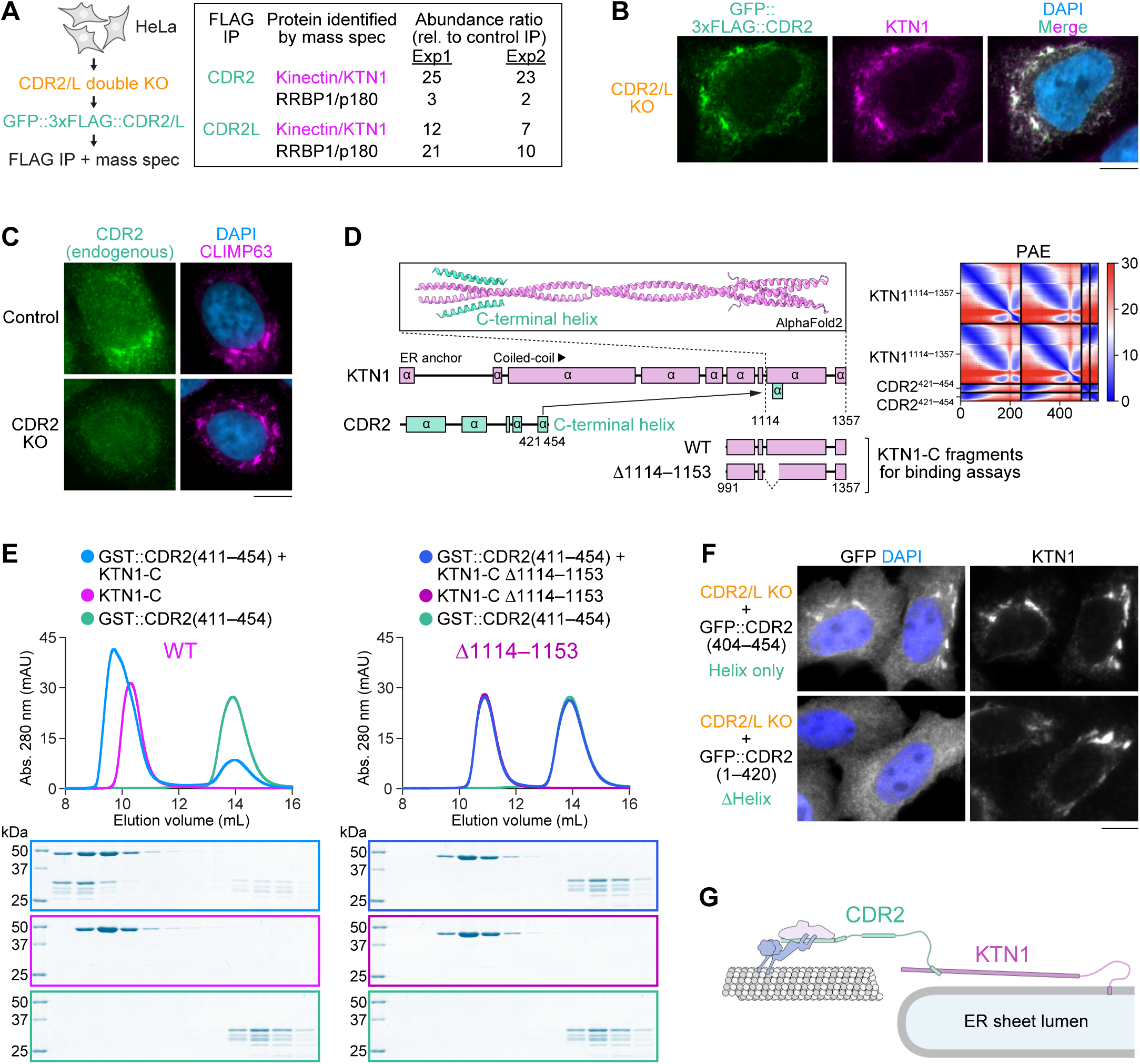
CDR2 and CDR2L interact and co-localize with the integral ER membrane protein KTN1. **(A)** Schematic illustrating construction of HeLa CDR2/L double KO cell lines stably expressing exogenous GFP::3xFLAG-tagged CDR2 or CDR2L used for immunoprecipitation followed by quantitative mass spectrometry. The relative abundance of KTN1 and RRBP1/p180 in anti-FLAG immunoprecipitations from transgenic and parental CDR2/L double KO cells is shown for two independent experiments (Exp1 and 2) on the right. **(B)** Immunofluorescence image of a HeLa cell stably expressing GFP::3xFLAG-tagged CDR2, showing co-localization with the ER sheet protein KTN1. Scale bar, 5 µm. **(C)** Immunofluorescence showing co-localization of endogenous CDR2 with the ER sheet protein CLIMP63. A CDR2 KO cell serves as the control for CDR2 antibody specificity. Scale bar, 5 µm. **(D)** AF2 model and predicted alignment error (PAE) plot of the KTN1 C-terminal coiled-coil domain in complex with the C-terminal helix of CDR2. KTN1 domain organization and C-terminal KTN1 fragments (KTN1-C) used for *in vitro* binding assays in *(E)* are also shown. **(E)** Elution profiles and BlueSafe-stained SDS-PAGE gels of purified recombinant human CDR2 and KTN1 fragments after SEC. The elution profile and gel for CDR2 are shown on both left and right to facilitate comparison between wild-type KTN1-C and the Δ1114–1153 mutant. Molecular weight is indicated in kilodaltons (kDa). **(F)** Immunofluorescence images of HeLa CDR2/L double KO cells transiently expressing GFP::CDR2 with and without its C-terminal helix, demonstrating that the helix is necessary and sufficient for ER localization. Scale bar, 5 µm. **(G)** Cartoon of the dynein recruitment pathway at ER sheets, based on results from *in vitro* reconstitution of protein–protein interactions and cell-based assays with binding-deficient mutants.

### A C-terminal helix in CDR2 is necessary and sufficient for binding to KTN1 and recruitment to ER sheets

To determine whether CDR2 and KTN1 directly bind each other, we first used AlphaFold2 (AF2) (Jumper *et al*., 2021) to predict interacting domains. KTN1 consists of an N-terminal transmembrane domain anchored in the ER membrane followed by a 1000-residue cytoplasmic domain that forms multiple segments of parallel dimeric coiled-coil. Structure prediction identified a high-confidence interaction between the last coiled-coil segment of KTN1 and the C-terminal helix in CDR2, which is highly conserved in CDR2 and CDR2L homologs from vertebrate and invertebrate species (Fig. 2D; Fig. S2D). SEC with purified recombinant KTN1(991–1357) and GST::CDR2(411–454) confirmed this interaction, which was abolished when the predicted CDR2 binding site in KTN1 was deleted (Δ1114–1153) (Fig. 2E). In CDR2/L double KO cells, the GFP-tagged C-terminal CDR2 helix (404–454) localized to ER sheets, whereas GFP::CDR2 lacking the C-terminal helix (1–420) did not (Fig. 2F). Interestingly, knockdown of KTN1 by RNAi not only delocalized CDR2 but also decreased its total levels (Fig. S2E–G), suggesting that the KTN1–CDR2 interaction stabilizes CDR2. We conclude that KTN1 recruits CDR2 to ER sheets through a direct interaction between their C-termini (Fig. 2G).

Given that the binding site for the C-terminal helix of CDR2 and CDR2L is conserved in the KTN1 paralog p180 (Fig. S2H), dynein is likely recruited to the ER by both of these integral membrane proteins. Interestingly, p180 was selectively enriched in immunoprecipitates of GFP::3xFLAG::CDR2L (Fig. 2A), suggesting that CDR2 and CDR2L may preferentially bind to KTN1 and p180, respectively.

### Double knockout of CDR2 and CDR2L promotes organization of ER sheets into stacks

To address the function of CDR2 and CDR2L, we examined ER morphology in CDR2/L double KO cells. Immunofluorescence showed that the naturally patchy distribution of the ER sheet components KTN1 and CLIMP63 became significantly more patchy in CDR2/L double KO cells (Fig. 3A; Fig. S3A). Correlative light–electron microscopy revealed that the bright µm-sized KTN1 patches observed by immunofluorescence correspond to stacks of ER sheets (Fig. S3B). In the absence of CDR2 and CDR2L, the fraction of cells with bright KTN1 patches was increased (Fig. 3A), and ER sheet stacks were larger (more sheets per stack), as determined by transmission electron microscopy (TEM) (Fig. 3B). Depleting KTN1 by RNAi in CDR2/L double KO cells essentially abolished ER sheet stacking but not formation of ER sheets *per se* (Fig. 3C, D; Fig. S3D), consistent with prior work implicating ER sheet proteins in stacking (Shibata *et al*., 2010). Immunoblotting showed that KTN1 levels were unchanged in CDR2/L double KO cells (Fig. S2G), suggesting enhanced stacking is not due to KTN1 overexpression. Instead, the brightness of KTN1 patches may indicate that KTN1 distribution within the ER becomes more concentrated on sheets in the absence of CDR2 and CDR2L. The CDR2/L double KO phenotype could be reversed by expressing exogenous wild-type GFP::CDR2 but not CDR2 lacking its CC1 box (Δ23– 39) or C-terminal helix (Δ421–454) (Fig. 3E–G; Fig. S3C). Collectively, these results suggest that CDR2 opposes the KTN1-dependent organization of ER sheets into stacks, and that this requires CDR2 recruitment by KTN1 and the interaction between CDR2 and DLIC (Fig. 3H).

**Figure 3:**
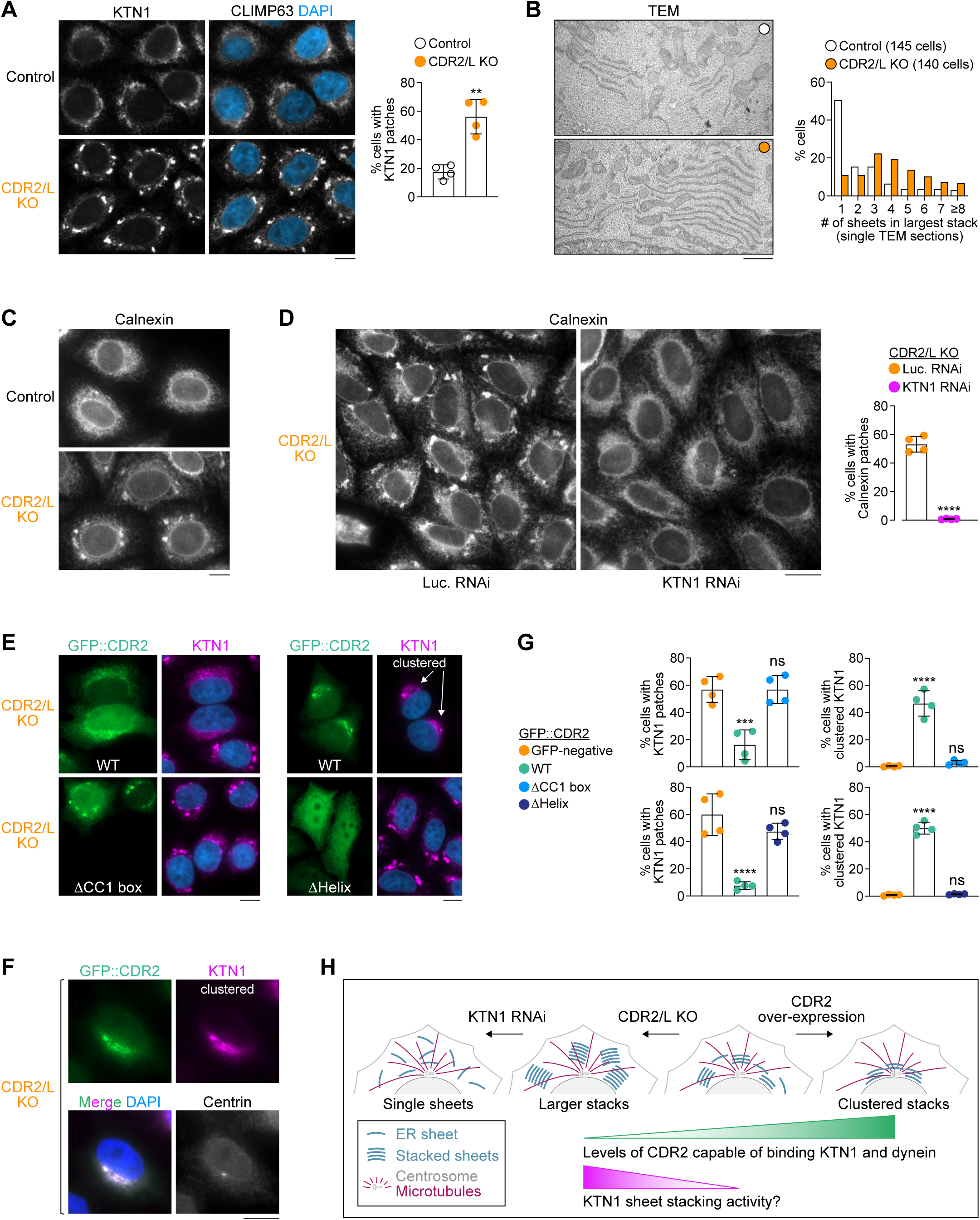
CDR2 regulates the organization of ER sheets. **(A)** *(left)* Immunofluorescence images showing exacerbated patchy distribution of KTN1 and CLIMP63 in HeLa CDR2/L double KO cells. Scale bar, 5 µm. *(right)* Fraction of cells with prominent KTN1 patches, plotted as mean ± SD (4 independent experiments, >1000 cells scored in total per condition). Statistical significance was determined using a two-tailed t test. \*\**P* < 0.01. See also *Fig. S3A*. **(B)** *(left)* Transmission electron microscopy (TEM) images of ER sheets in control and CDR2/L double KO cells, both treated with siRNAs against Luciferase to facilitate comparison with KTN1 depletion in *Fig. S3D*. Scale bar, 1 µm. *(right)* Number of ER sheets present in the largest stack identified in individual cells using single TEM sections. The total number of cells analyzed in 3 independent experiments is indicated. **(C), (D)** Immunofluorescence showing patchy distribution of the ER protein Calnexin in CDR2/L double KO cells, which is abolished after knockdown of KTN1 by RNA interference (RNAi). Luciferase (Luc.) RNAi serves as the control. Scale bars, 10 µm *(C)* and 20 µm *(D)*. The fraction of cells with prominent Calnexin patches is plotted as mean ± SD (4 independent experiments, >1500 cells scored in total per condition). Statistical significance was determined using a two-tailed t test. \*\*\*\**P* < 0.0001. **(E), (F)** Immunofluorescence images of CDR2/L double KO cells transiently transfected with WT GFP::CDR2 or mutants lacking residues 23–39 (ΔCC1 box) or 404–454 (ΔHelix). Centrin-3 staining in *(F)* shows that WT GFP::CDR2 and KTN1 cluster together at centrosomes. Images in *(E)* include examples of untransfected cells (GFP-negative) for comparison. Scale bars, 10 µm. **(G)** Fraction of cells (mean ± SD, 4 independent experiments, >600 cells scored in total per condition) with prominent KTN1 patches *(left)* and centrosome-proximal KTN1 clustering *(right)* in the conditions shown in *(E)*. ΔCC1 box and ΔHelix experiments each have their own WT and GFP-negative controls. Statistical significance was determined using ordinary one-way ANOVA followed by Tukey’s multiple comparisons test. \*\*\*\**P* < 0.0001; \*\*\**P* < 0.001; *ns* = not significant, *P* > 0.05. **(H)** Cartoon summarizing the effect of CDR2 and KTN1 levels on ER sheet organization.

### CDR2 overexpression results in CC1 box-dependent clustering of ER sheets near centrosomes

Rescue experiments with exogenous GFP::CDR2 in CDR2/L double KO cells revealed that in addition to reversing excessive ER sheet stacking, GFP::CDR2 frequently induced clustering of ER sheets near centrosomes, marked by centrin-3 staining (Fig. 3E–G). By contrast, ER sheet clustering in CDR2/L double KO cells was never observed with exogenous GFP::CDR2 lacking the CC1 box or C-terminal helix. ER sheets also did not cluster appreciably in untransfected control cells, suggesting that clustering is specifically induced by exogenous GFP::CDR2. These results support the idea that KTN1-associated CDR2 can recruit dynein activity to promote centrosome-directed transport of ER sheets (Fig. 3H).

To compare CDR2’s ability to recruit dynein activity to that of another established activating adaptor, we replaced the CDR2 N-terminal region (1–185) with that of JIP3 (Singh *et al*., 2024). The JIP3(1–185)::CDR2(186–454) chimera co-localized with KTN1 in CDR2/L double KO cells and induced penetrant and tight clustering (Fig. S3E). This indicates that the CDR2 N-terminus is less efficient than that of JIP3 at dynein recruitment and/or activation at ER sheets, which would be consistent with our results from *in vitro* motility assays. Alternatively, the efficiency of the JIP3::CDR2 chimera may reflect the absence of autoinhibition mechanisms present in full-length CDR2.

### CDR2 competes with eEF1Bβ, but not KIF5C, for binding to KTN1

Prior studies identified the kinesin-1 heavy chain KIF5 and the eEF1Bβ subunit of the translation elongation factor 1 complex (eEF1) as direct binding partners of the KTN1 C-terminus (Ong *et al*., 2000, 2003, 2006). The KIF5 binding site was mapped to KTN1 residues 1188-1288 (Ong *et al*., 2000). CDR2 and KIF5 therefore occupy adjacent, non-overlapping sites. By contrast, structure prediction suggested that a helix formed by eEF1Bβ residues 33–60 occupies the same site on KTN1 as the CDR2 helix (Fig. 4A; Fig. S3F). SEC with purified recombinant proteins demonstrated that GST::eEF1Bβ(30–66) forms a robust complex with KTN1(991–1357), but not when the CDR2 binding site in KTN1 is deleted (Δ1114–1157) (Fig. 4B). Using GST pull-downs, we confirmed that KTN1(991–1357) binds GST::KIF5C(807–956) and showed that this interaction is insensitive to deletion of the CDR2/eEF1Bβ binding site (Fig. 4C). In cells, eEF1Bβ co-localized with KTN1 and CDR2 (Fig. 4D, E), and replacing the CDR2 helix with the eEF1Bβ helix was sufficient to localize the GFP-tagged chimera to ER sheets (Fig. 4F). Furthermore, overexpression of GFP::CDR2 lacking its CC1 box (Δ23–39) displaced eEF1Bβ from KTN1 patches in CDR2/L double KO cells (Fig. 4G). We conclude that eEF1Bβ, but not KIF5, competes with CDR2 for binding to KTN1 and recruitment to ER sheets.

**Figure 4:**
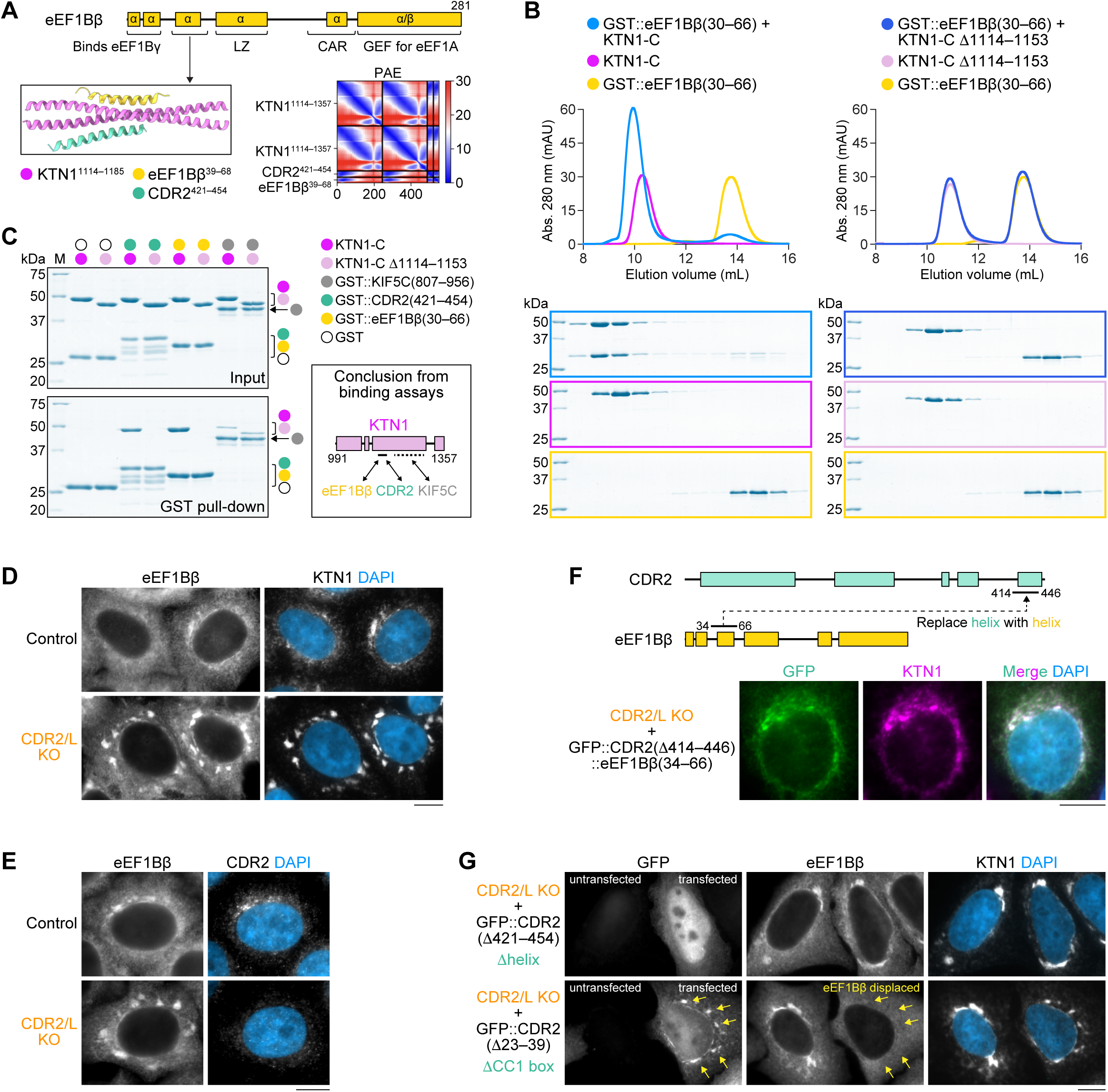
CDR2 competes with eEF1BΔ, but not KIF5, for binding to KTN1 and localization to ER sheets. **(A)** Domain organization of the β subunit of eukaryotic translation elongation factor 1B (eEF1Bβ, UniProt P29692; encoded by gene EEF1D). AF2 model and predicted alignment error (PAE) plot shows that an N-terminal eEF1Bβ helix and the C-terminal CDR2 helix occupy the same site in KTN1. See *Fig. S3F* for a prediction of full-length eEF1Bβ in complex with KTN1. **(B)** Elution profiles BlueSafe-stained SDS-PAGE gels of purified recombinant human eEF1Bβ and KTN1 fragments after SEC. The elution profile and gel for eEF1Bβ are shown on both left and right to facilitate comparison between wild-type KTN1-C and the Δ1114–1153 mutant. Molecular weight is indicated in kilodaltons (kDa). **(C)** BlueSafe-stained SDS-PAGE gels of purified recombinant proteins prior to addition of glutathione agarose resin (Input) and after elution from the resin (GST pull-down), showing that the binding site of KIF5C on KTN1 is distinct from that of CDR2/eEF1Bβ. Schematic summarizes the results of binding assays. Dotted line indicates the KIF5C binding site on KTN1 mapped by Ong *et al*. (2000). **(D), (E)** Immunofluorescence demonstrating co-localization of eEF1Bβ, KTN1, and CDR2 in HeLa cells. Scale bars, 10 µm. **(F)** Domain swapping experiment showing that replacing the C-terminal CDR2 helix with the N-terminal eEF1Bβ helix (both 33 residues long) is sufficient to target GFP::CDR2 to the ER. Scale bar, 10 µm. **(G)** Immunofluorescence showing that overexpression of GFP::CDR2 displaces eEF1Bβ from the ER (arrows), but only if CDR2 can bind KTN1. Scale bar, 10 µm.

### eEF1Bβ knockdown enhances recruitment of endogenous CDR2 to ER sheets and promotes ER sheet clustering near centrosomes

Given that eEF1Bβ is an abundant protein, eEF1Bβ levels may be limiting for CDR2 recruitment to ER sheets due to competition for KTN1 binding. To test this idea, we decreased eEF1Bβ levels by RNAi. Immunoblotting showed that overall levels of CDR2 slightly increased (∼1.4 fold) and KTN1 levels remained unchanged after eEF1Bβ knockdown (Fig. 5A). CDR2 localization to ER sheets was significantly more pronounced in eEF1Bβ-depleted cells, consistent with the idea that KTN1 is now free to bind CDR2 (Fig. 5B). Strikingly, ER sheets with elevated CDR2 levels tended to cluster near centrosomes (Fig. 5C, D). By contrast, ER sheet distribution remained unchanged when eEF1Bβ was knocked down in CDR2/L double KO cells (Fig. 5C; Fig. S3G). These results support the idea that enhanced recruitment of CDR2 to ER sheets promotes ER sheet clustering near centrosomes. Taken together, our findings suggest that competitive binding of the dynein adaptor CDR2 and eEF1Bβ to KTN1 regulates ER sheet organization.

**Figure 5:**
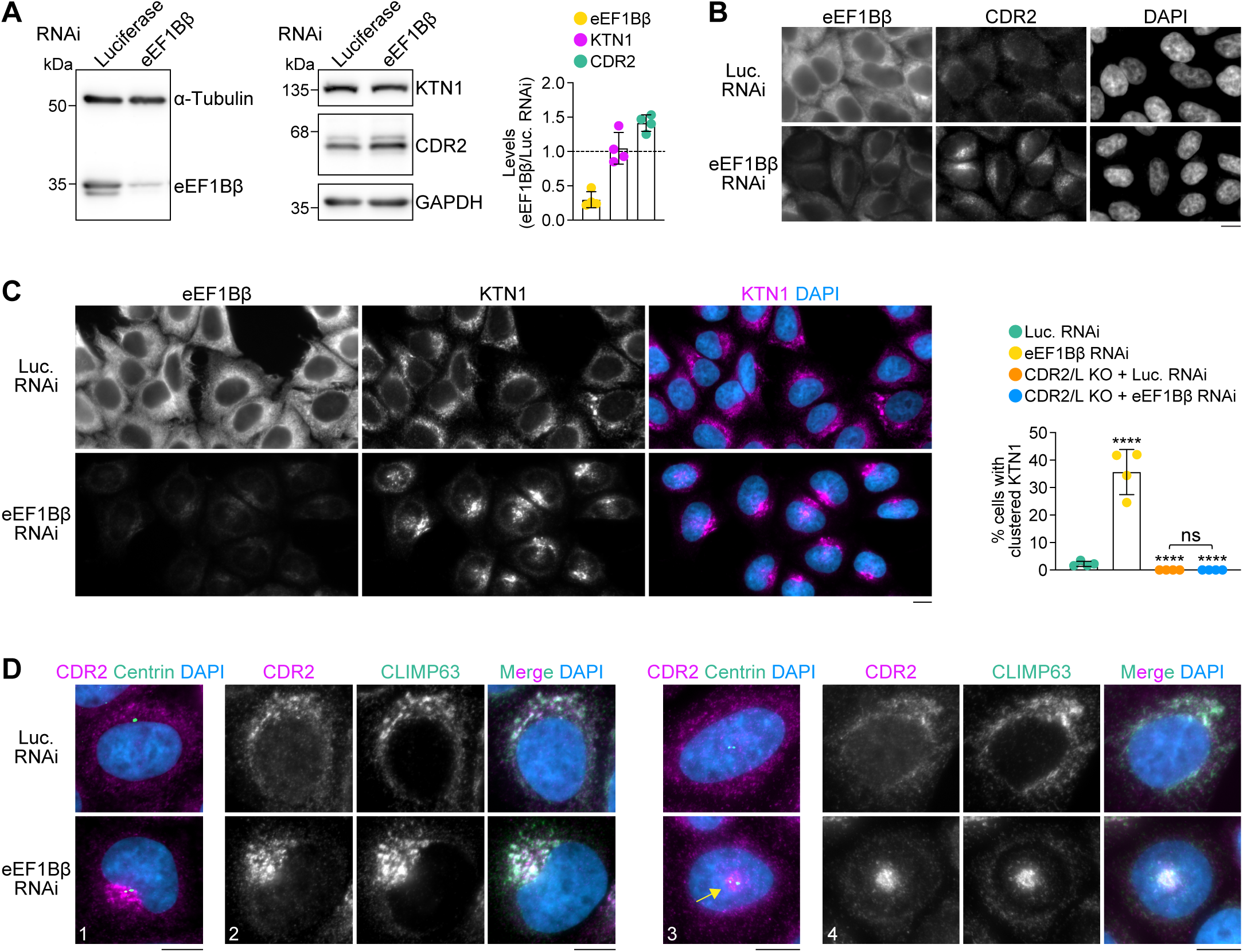
eEF1Bβ knockdown enhances CDR2 recruitment to ER sheets and promotes ER sheet clustering near centrosomes. **(A)** Immunoblots and quantification of protein levels in HeLa cells treated with siRNAs against Luciferase or eEF1Bβ. The 3 immunoblots on the right are from the same membrane. Protein levels relative to Luciferase RNAi, quantified based on immunoblot signal intensity after normalization to the loading control (α-tubulin for eEF1Bβ, GAPDH for CDR2 and KTN1), are plotted as mean ± SD (4 independent experiments). **(B)** Immunofluorescence images showing enhanced ER localization of CDR2 in eEF1Bβ-depleted HeLa cells. Scale bar, 10 µm. **(C)** Immunofluorescence images (maximum intensity projection of z-stack) showing that eEF1Bβ knockdown results in redistribution of KTN1 into clusters. See *Fig. S3G* for corresponding immunofluorescence images in CDR2/L double KO cells. The fraction of mock-and eEF1Bβ-depleted cells with a clustered KTN1 distribution is plotted as mean ± SD (4 independent experiments, >1300 cells scored in total per condition). Statistical significance was determined using ordinary one-way ANOVA followed by Tukey’s multiple comparisons test. \*\*\*\**P* < 0.0001; *ns* = not significant, *P* > 0.05. **(D)** Immunofluorescence images (maximum intensity projection of z-stack) illustrating that CDR2 and CLIMP63 clustering together at centrosomes in eEF1Bβ-depleted cells. Four examples are shown, in which the ER clusters either on the side (cells 1 and 2) and on top (cells 3 and 4) of the nucleus. Scale bars, 10 µm.

### Conclusions

Here we identify and dissect a molecular pathway that recruits dynein to ER sheets and regulates ER organization in a human cancer cell line. The pathway involves the novel dynein adaptors CDR2 and CDR2L, which use a conserved C-terminal helix to bind an equally well conserved site on the related integral ER membrane proteins KTN1 and p180. We show that these interactions place dynein in proximity to kinesin-1 at the C-terminus of KTN1/p180’s extended cytosolic coiled-coil domain. This is in line with the emerging realization that dynein is often paired on cargo with a kinesin (Abid Ali *et al*., 2023, *Preprint*; Canty *et al*., 2023; Celestino *et al*., 2022; Cmentowski *et al*., 2023; Kendrick *et al*., 2019; Splinter *et al*., 2010) and implies close collaboration between the two motors at ER membranes.

One role for KTN1-associated dynein may be to oppose kinesin-1-mediated transport of KTN1 to the cell periphery, which has been functionally linked to the maturation of focal adhesions (Guadagno *et al*., 2020; Ng *et al*., 2016; Santama *et al*., 2004; Zhang *et al*., 2010). Our finding that CDR2 overexpression induces perinuclear clustering of KTN1 is consistent with this idea. However, contrary to what would be expected from such a role, knocking out CDR2 and CDR2L does not result in KTN1 accumulation at the cell periphery. Instead, double knockout cells have an altered ER organization characterized by enlarged ER sheet stacks enriched in KTN1. ER sheets are specialized in protein translocation, and it is envisioned that membrane-bound polysomes cooperate with the sheet-enriched membrane proteins KTN1, p180 and CLIMP63 to form segregated rough ER domains in mammalian cells (Shibata *et al*., 2006, 2010). This involves the concentration of sheet-enriched proteins by polysomes and vice-vera, which in turn is expected to promote sheet stacking (Shibata *et al*., 2010; Terasaki *et al*., 2013). If transport of KTN1 along microtubules via CDR2/CDR2L–dynein opposes its concentration on sheets, it would explain why loss of CDR2 and CDR2L enhances sheet stacking. Our finding that CDR2 competes for recruitment to KTN1 with eEF1Bβ, which anchors the eEF1 complex at ER sheets (Ong *et al*., 2003, 2006), additionally suggests that the absence of CDR2 and CDR2L may promote sheet stacking by reinforcing the association of KTN1 with polysomes via eEF1. Whether and how KTN1-associated kinesin-1 affects this process, and, more broadly, how motor recruitment to the C-terminus of KTN1 functionally relates to its N-terminal microtubule-binding activity, recently reported to control ER distribution (Zheng *et al*., 2022), are interesting questions for the future. Taken together, our results support the idea of an overall antagonistic relationship between dynein-driven ER dynamics mediated by CDR2/CDR2L and protein biosynthesis at ER membranes. Intriguingly, the recent identification of *D. melanogaster* Centrocortin as a dynein adaptor that transports its mRNA to centrosomes (Zein-Sabatto *et al*., 2024, *Preprint*) hints at the possibility that CDR2 and CDR2L could facilitate their own translation at the ER.

CDR2 and CDR2L are prominently expressed in the mammalian brain (Hwang *et al*., 2016; Raspotnig *et al*., 2022). Neurons, characterized by uniquely compartmentalized ER organization (Farías *et al*., 2019; Koppers *et al*., 2024; Renvoisé and Blackstone, 2010), may therefore offer a relevant physiological context in which to further explore the roles of CDR2 and CDR2L in ER organization. In patients with PCD, the predominant tumor-induced autoantibody present in serum and cerebrospinal fluid is anti-Yo, which recognizes CDR2 and CDR2L (Kråkenes *et al*., 2019; Sakai *et al*., 1990). Anti-Yo can be taken up by Purkinje cells *in vivo* (Graus *et al*., 1991; Greenlee *et al*., 1995), and the intracellular interaction between anti-Yo and CDR proteins induces Purkinje cell death *in vitro* (Greenlee *et al*., 2010; Schubert *et al*., 2014). Our study raises the possibility that the toxicity following anti-Yo uptake stems from pathologic changes in ER organization, a factor implicated in various neurological disorders (Perkins and Allan, 2021; Westrate *et al*., 2015). EM studies with anti-Yo antibody have indeed demonstrated reactivity with ER-associated antigens (Hida *et al*., 1994; Rodriguez *et al*., 1988), and disrupted ER function caused by anti-Yo exposure would be consistent with reports that anti-Yo impairs calcium homeostasis in Purkinje cells (Panja *et al*., 2019: Schubert *et al*., 2024). Our findings may therefore open the door to a better understanding of PCD pathogenesis.

## MATERIALS AND METHODS

### DNA constructs

#### Tissue culture

For transient expression of CDR2, CDR2L, eEF1Bβ and JIP3, cDNA was inserted into a pcDNA5-FRT-TO-based vector (Invitrogen) modified to contain N-terminal Myc::EGFP::TEV::S-peptide. To generate cell lines stably expressing GFP::3xFLAG-tagged CDR2 and CDR2L, cDNA for 3xFLAG::CDR2 and 3xFLAG::CDR2L was inserted into pLenti-CMV-GFP-Hygro (Addgene 17446). To generate CDR2/CDR2L single and double KO cells by CRISPR/Cas9, protospacer sequences targeting CDR2 (GCTGGCGGAAAACCTGGTAG; CTACCAGGTTTTCCGCCAGC; ACAATTAGACGTCACAGCAA; TTGCTGTGACGTCTAATTGT) and CDR2L (GCTGGTCGTACCAGGACTCC; CTGGTACGACCAGCAGGACC) were inserted into the BsmBI sites of pLenti-sgRNA (Addgene 71409) or pKM808 (Addgene 134181).

#### Biochemistry

For expression in insect cells, we used previously described full length human cytoplasmic dynein-1 with a C-terminal ZZ-SNAPf tag on DHC (Schlager *et al*., 2014) and human Lis1 with an N-terminal ZZ-TEV tag (Baumbach *et al*., 2017). The DHC construct contained mutations in the linker (R1567E and K1610E) to help overcome the autoinhibited conformation (Zhang *et al*., 2017). For bacterial expression of CDR2, CDR2L, DLIC1, eEF1BΔ, JIP3, KIF5 and KTN1 fragments, cDNA was inserted into a 2CT vector containing an N-terminal 6xHis::maltose binding protein (MBP) followed by a TEV protease cleavage site and C-terminal Strep-tag II, or into pGEX-6P-1 containing N-terminal glutathione S-transferase (GST) followed by a Prescission protease cleavage site and C-terminal 6xHis.

Protein residue numbers in text and figures refer to the following UniProt entries Q01850 (CDR2_HUMAN), Q86X02 (CDR2L_HUMAN), Q9Y6G9 (DC1L1_HUMAN), Q14204 (DYHC1_HUMAN), P29692 (EF1D_HUMAN), Q9UPT6 (JIP3_HUMAN), P28738 (KIF5C_MOUSE) and Q86UP2 (KTN1_HUMAN).

### Protein expression

Cytoplasmic dynein-1 and Lis1 were expressed using the Sf9/baculovirus system. Fresh bacmid DNA was transfected into Sf9 cells at 0.5×10^6^ cells/mL in 6-well cell culture plates using FuGene HD (Promega) according to the manufacturer’s protocol (final concentration 10 µg/mL). After six days, 1 mL of the culture supernatant was added to 50 mL of 1×10^6^ cells/mL and cells were infected for five days in a shaking incubator at 27°C. P2 virus was isolated by collecting the supernatant after centrifugation at 4,000 rcf for 15 min and stored at 4°C. For expression, 10 mL of P2 virus was used to infect 1 L of Sf9 cells at 1.5-2×10^6^ cells/mL for 72 hours in a shaking incubator at 27°C. Cells were harvested by centrifugation at 4,000 rcf for 10 min at 4°C, and washed with cold PBS. The cell pellet was flash frozen and stored at -80°C.

For expression of CDR2, CDR2L, DLIC1, eEF1BΔ, JIP3, KIF5 and KTN1 fragments, plasmids were transformed into the *E. coli* Rosetta strain. Single colonies were grown in 10 mL Luria-Bertani medium overnight at 37°C in a shaking incubator. 10 mL of saturated culture were diluted into 1 L and incubated at 30°C until an OD_600_ of 0.5. Expression was induced with 0.1 mM IPTG and cultures were grown overnight at 18°C. Cells were harvested by centrifugation at 4,000 rcf for 15 min at 4°C in a Mega Star 4.0R centrifuge with TX-1000 rotor (Avantor), and cell pellets were stored at -80°C.

### Protein purification

Dynactin was purified from frozen porcine brains as previously described (Urnavicius *et al*. 2015). Fresh brains were cleaned in homogenization buffer (35 mM PIPES pH 7.2, 5 mM MgSO_4_, 100 µM EGTA, 50 µM EDTA), and flash frozen in liquid nitrogen. Frozen brains were broken into pieces using a hammer. The brain pieces were blended and resuspended in homogenization buffer supplemented with 1.6 mM PMSF, 1 mM DTT, and 4 cOmplete EDTA-free protease inhibitor cocktail tablets (Roche) per 500 mL. After thawing, the lysate was centrifuged in a JLA 16.250 rotor (Beckman Coulter) at 16,000 rpm for 15 min at 4°C. The supernatant was further clarified in a Type 45 Ti rotor (Beckman Coulter) at 45,000 rpm for 50 min at 4°C. After filtering the supernatant in a Glass Fiber filter (Sartorius) and a 0.45 µm filter (Elkay Labs), it was loaded on a column packed with 250 mL of SP-Sepharose (Cytiva) pre-equilibrated with SP buffer (35 mM PIPES pH 7.2, 5 mM MgSO_4_, 1 mM EGTA, 0.5 mM EDTA, 1 mM DTT, 0.1 mM ATP) using an ÄKTA Pure system (Cytiva). The column was washed with SP buffer with 3 mM KCl before being eluted in a linear gradient up to 250 mM KCl over 3 column volumes. The peak around ∼15 mS/cm was collected and filtered with a 0.22 µm filter (Elkay Labs) before being loaded on a MonoQ 16/10 column (Cytiva) pre-equilibrated with MonoQ buffer (35 mM PIPES pH 7.2, 5 mM MgSO_4_, 100 µM EGTA, 50 µM EDTA, 1 mM DTT). The column was washed with MonoQ buffer before being eluted in a linear gradient up to 150 mM KCl over 1 column volume, followed by another linear gradient up to 350 mM KCl over 10 column volumes. The peak around ∼39 mS/cm was pooled and concentrated to ∼3 mg/mL before being loaded on a TSKgel G4000SWXL column (Tosoh Bioscience) preequilibrated with GF150 buffer (25 mM HEPES pH 7.2, 150 mM KCl, 1 mM MgCl_2_) supplemented with 5 mM DTT and 0.1 mM ATP. The peak at ∼114 mL was pooled and concentrated to ∼3 mg/mL. 3 µL aliquots were flash frozen in liquid nitrogen and stored at -80°C.

For dynein purification, a cell pellet from 1 L expression was resuspended in 50 mL lysis buffer (50 mM HEPES pH 7.4, 100 mM NaCl, 10% (v/v) glycerol, 0.1 mM ATP) supplemented with 2 mM PMSF, 1 mM DTT, and 1 cOmplete EDTA-free protease inhibitor cocktail tablet. Cells were lysed using a 40-mL dounce tissue grinder (Wheaton) with ∼20 strokes. The lysate was clarified at 503,000 rcf for 45 min at 4°C using a Type 70 Ti rotor (Beckman Coulter). The supernatant was incubated with 3 mL IgG Sepharose 6 Fast Flow beads (Cytiva) pre-equilibrated with lysis buffer for 4 hours at 4°C. The beads were applied to a gravity flow column and washed with 150 mL of lysis buffer and 150 mL of TEV buffer (50 mM Tris-HCl pH 7.4, 150 mM KAc, 2 mM MgAc, 1 mM EGTA, 10% (v/v) glycerol, 0.1 mM ATP, 1 mM DTT). For TMR labeled dynein, beads were transferred to a tube and incubated with 10 μM SNAP-Cell TMR-Star dye (New England Biolabs) for 1 hour at 4°C prior to the TEV buffer washing step. The beads were then transferred to a 5-mL centrifuge tube (Eppendorf) and filled up completely with TEV buffer. 400 μg TEV protease was added to the beads followed by overnight incubation at 4°C. The beads were transferred to a gravity flow column and the flow through containing the cleaved protein was collected. The protein was concentrated to ∼2 mg/mL and loaded onto a TSKgel G4000SWXL column pre-equilibrated with GF150 buffer supplemented with 5 mM DTT and 0.1 mM ATP. Peak fractions were pooled and concentrated to ∼2.5–3 mg/mL. Glycerol was added to a final concentration of 10% from an 80% stock made in GF150 buffer. 3 µL aliquots were flash frozen and stored at -80°C.

For Lis1 purification, a cell pellet from 1 L expression was resuspended in 50 mL lysis buffer B (50 mM Tris-HCl pH 8, 250 mM KAc, 2 mM MgAc, 1 mM EGTA, 10% (v/v) glycerol, 0.1 mM ATP, 1 mM DTT) supplemented with 2 mM PMSF. Cells were lysed using a 40-mL dounce tissue grinder (Wheaton) with ∼20 strokes. The lysate was clarified at 150,000 rcf for 30 min at 4°C using a Type 45 Ti rotor (Beckman Coulter). The supernatant was incubated with 3 mL IgG Sepharose 6 Fast Flow beads (Cytiva) pre-equilibrated with lysis buffer B for 4 hours at 4°C. The beads were then applied to a gravity flow column and washed with 150 mL of lysis buffer B. The beads were then transferred to a 5-mL centrifuge tube (Eppendorf) and filled up completely with lysis buffer B. 400 μg TEV protease was added to the beads followed by overnight incubation at 4°C. The beads were transferred to a gravity flow column and the flow through containing the cleaved protein was collected. The protein was concentrated to ∼5 mg/mL and loaded onto a Superdex 200 Increase 10/300 column (Cytiva) pre-equilibrated with GF150 buffer supplemented with 5 mM DTT and 0.1 mM ATP. Peak fractions were pooled and concentrated to ∼5 mg/mL. Glycerol was added to a final concentration of 10% from an 80% stock made in GF150 buffer. 5 µL aliquots were flash frozen and stored at -80°C.

For purification of CDR2, CDR2L, JIP3, and KTN1 fragments with a C-terminal Strep-tag II (StTgII), bacterial pellets from 1 L expression were resuspended in 30 mL lysis buffer C (20 mM Tris-HCl pH 8, 300 mM NaCl, 10 mM imidazole, 1 mM DTT) supplemented with 1 mM PMSF and 2 mM benzamidine-HCl. Cells were lysed with a cell cracker, and the lysate was cleared twice by centrifugation at 40,000 rcf for 20 min each at 4°C using a JA 25.50 rotor (Beckman Coulter). The cleared lysate was incubated with 2 mL Ni-NTA resin (Thermo Fisher Scientific) for 1 hour at 4 °C, transferred to a gravity flow column (Pierce), and washed with 150 mL wash buffer (20 mM Tris-HCl pH 8, 300 mM NaCl, 20 mM imidazole, 0.1% (v/v) Tween 20, 1 mM DTT) supplemented with 2 mM benzamidine-HCl. Proteins were eluted with 10 mL elution buffer (20 mM Tris-HCl pH 8, 300 mM NaCl, 250 mM imidazole, 1 mM DTT) and incubated overnight at 4°C with 130 µg TEV protease. Cleaved proteins were incubated with 2 mL Strep-Tactin Sepharose resin (IBA) for 1 hour at 4°C, transferred to a gravity flow column, washed with 2 × 50 mL wash buffer B (20 mM Tris-HCl pH 8, 300 mM NaCl) and 50 mL wash buffer C (20 mM Tris-HCl pH 8, 150 mM NaCl), and eluted with 10 mL elution buffer B (100 mM Tris-HCl pH 8, 150 mM NaCl, 10 mM *d*-desthiobiotin). Proteins were concentrated and loaded onto a Superdex 200 Increase 10/300 GL column pre-equilibrated with storage buffer (25 mM HEPES pH 7.5, 150 mM NaCl, 1mM DTT). Peak fractions were pooled and concentrated, glycerol was added to 10% (v/v), and aliquots were flash frozen in liquid nitrogen and stored at -80°C.

For purification of CDR2, DLIC1, eEF1Bβ, and KIF5C fragments with an N-terminal GST and C-terminal 6xHis tag, bacterial pellets from 1 L expression were lysed and proteins were purified with Ni-NTA resin, as described above. After elution from Ni-NTA resin, proteins were incubated with 2 mL Pierce Glutathione Agarose resin (Thermo Fisher Scientific) for 1 hour at 4°C, transferred to a gravity flow column, washed with 2 × 50 mL wash buffer B and 50 mL wash buffer C, and eluted with 10 mL elution buffer C (25 mM HEPES pH 7.5, 150 mM NaCl, 10 mM reduced L-glutathione). For KIF5C, L-glutathione was removed with an Econo-Pac 10DG column (Bio-Rad) by buffer exchange into storage buffer, and the protein was concentrated, flash frozen and stored at -80°C, as described for StTgII proteins. For CDR2, DLIC1, and eEF1BΔ, proteins were concentrated and loaded onto a Superdex 200 Increase 10/300 GL column pre-equilibrated with storage buffer. Peak fractions were pooled, concentrated, flash frozen and stored at -80°C, as described for StTgII proteins.

### GST pull-downs

Proteins (250 pmol each) were mixed in a total of 20 µL PD buffer (25 mM HEPES pH 7.5, 150 mM NaCl, 5 mM DTT) in a 1.5-mL tube and incubated at room temperature for 30 min. 4 µL were removed from the mixture and added to a tube containing 23 µL PD buffer and 9 µL 4× SDS-PAGE sample buffer (200 mM Tris-HCl pH 6.8, 40% (v/v) glycerol, 8% (w/v) SDS, 400 mM DTT, 0.4% (w/v) bromophenol blue) ("Input"). To the remaining 16 µL protein mixture, 30 µL of a 50% slurry of Pierce Glutathione Agarose resin/PD buffer were added and the resin/protein mixture was rotated horizontally for 30 min. The resin was washed quickly with 3 × 500 µL PD buffer using 15-s spins in a microfuge (Roth), all buffer was removed with a gel loading tip, and the resin was incubated with 50 µL elution buffer C for 15 min with rotation. The resin was pelleted at 20,000 rcf for 1 min in an Eppendorf 5424 centrifuge, 36 µL of the eluate were removed and added to a tube containing 12 µL 4× SDS-PAGE sample buffer, and the sample was heated for 1 min at 95°C ("GST pull-down"). 7 µL of "Input" and "GST pull-down" samples were separated by 14% SDS-PAGE and proteins were visualized with BlueSafe stain (NZY Tech).

### Strep-Tag II pull-downs from porcine brain lysate

Porcine brain lysate was prepared as described previously (McKenney *et al*., 2014). In brief, fresh brains were broken into small chunks, flash-frozen in liquid nitrogen, and stored at -80°C. Frozen brain chunks were homogenized in equal weight/volume of buffer (50 mM HEPES pH 7, 50 mM PIPES, 1 mM EDTA, and 2 mM MgSO_4_, 1 mM DTT) using a waring blender, followed by glass pestle grinding. After clarification at 34,000 rcf for 45 min at 4°C using a JA 25.50 rotor, the crude homogenate was flash frozen in 1-mL aliquots and stored at -80°C.

For pull-downs, 250 pmol of purified protein were added to 15 µL Strep-Tactin Sepharose resin pre-equilibrated in 100 µL PD buffer B (30 mM HEPES pH 7.5, 50 mM KAc, 2 mM Mg(Ac)_2_, 1 mM EGTA, 10% (v/v) glycerol, 0.1% (v/v) NP-40, 5 mM DTT) in a 1.5-mL tube and incubated for 30 min at room temperature. Porcine brain lysate was thawed, supplemented with 1 mM PMSF, and cleared at 20,000 rcf for 10 min at 4°C in a Megafuge 8R (Eppendorf). 1 µL was removed from the cleared lysate, added to a tube containing 39 µL 1× SDS-PAGE sample buffer, and heated for 3 min at 95°C ("Brain lysate"). 300 µL PD buffer B and 100 µL brain lysate were added to the protein/resin mixture (total volume ∼500 µL) and incubated with rotation for 1 hour at 4°C. The resin was washed quickly with 3 × 500 µL ice-cold PD buffer B using 15-s spins in a microfuge, all buffer was removed with a gel loading tip, and the resin was incubated with 50 µL elution buffer B for 15 min at room temperature with rotation. The resin was pelleted at 20,000 rcf for 1 min in an Eppendorf 5424 centrifuge, 36 µL of the eluate were removed, added to a tube containing 12 µL 4× SDS-PAGE sample buffer, and the sample was heated for 1 min at 95°C ("StTgII pull-down"). 8 µL of "Brain lysate" and "StTgII pull-down" samples were separated by 12% SDS-PAGE and visualized with BlueSafe stain, and 10 µL were separated by 12% SDS-PAGE and processed for immunoblotting, as described below.

### Size exclusion chromatography to assess protein complex formation

4 nmol (Fig. 2E; Fig. 4B) or 8 nmol (Fig. 1B; Fig. S1C) of each protein were diluted with storage buffer to a final volume of 200 µL in a 1.5-mL tube, corresponding a final concentration of 20 µM and 40 µM, respectively. After incubation at room temperature for 30 min, samples were cleared in a Megafuge 8R at 20,000 rcf for 10 min at 4°C and loaded onto a Superdex 200 Increase 10/300 GL column. SEC was performed at room temperature on an ÄKTA Pure 25L1 system at a flow rate of 0.5 mL/min. 0.5-mL fractions were collected and protein elution was monitored at 280 nm. 30 µL were removed from each fraction and added to a tube containing 10 µL 4× SDS-PAGE sample buffer. Samples were heated for 1 min at 95°C, and 5 µL were separated by 14% SDS-PAGE. Proteins were visualized by BlueSafe staining.

### In vitro TIRF motility assays

In vitro TIRF assays were carried out as previously described (Urnavicius *et al*., 2018). Microtubules were prepared the day before the assay was performed. Microtubules were made by mixing 1.5 μL of 2 mg/mL HiLyte Fluor 488 tubulin (Cytoskeleton), 2 μL of 2 mg/mL biotinylated tubulin (Cytoskeleton) and 6.5 μL of 13 mg/mL unlabelled pig tubulin (Schlager *et al*., 2014) in BRB80 buffer (80 mM PIPES pH 6.8, 1 mM MgCl2, 1 mM EGTA, 1 mM DTT). 10 μL of polymerization buffer (2× BRB80 buffer, 20% (v/v) DMSO, 2 mM Mg-GTP) was added followed by incubation for 5 min at 4°C. Microtubules were polymerized for 1 hour at 37°C. The sample was diluted with 100 μL MT buffer (BRB80 supplemented with 20 μM paclitaxel), then centrifuged on a benchtop centrifuge (Eppendorf) at 21,000 rcf for 9 min at room temperature. The resulting pellet was gently resuspended in 100 μL MT buffer, then centrifuged again as above. 50 μL MT buffer was added and the microtubule solution was protected from light. Before usage, and every 5 hours during data collection, the microtubule solution was spun again at 21,000 rcf for 9 min and the pellet resuspended in the equivalent amount of MT buffer.

Motility chambers were prepared by applying two strips of double-sided tape approximately 5 mm apart on a glass slide and then placing the coverslip on top. Before use, coverslips were pretreated by sequentially sonicating in 3 M KOH, water, and 100% ethanol followed by plasma treatment in an Ar:O_2_ (3:1) gas mixture for 3 min. Coverslips were functionalized using PLL-PEG-Biotin (SuSOS AG), washed with 50 μL TIRF buffer (30 mM HEPES pH 7.2, 5 MgSO_4_, 1 mM EGTA, 2 mM DTT) and incubated with 1 mg/mL streptavidin (New England Biolabs). The chamber was again washed with TIRF buffer and incubated with 10 μL of a fresh dilution of microtubules (1.5 μL microtubules diluted into 10 μL TIRF-Casein buffer [TIRF buffer supplemented with 50 mM KCl, 1 mg/mL casein and 5 µM paclitaxel]) for 1 min. Chambers were then blocked with 50 μL TIRF-Casein buffer.

Complexes were prepared by mixing each component in a total volume of 6 μL in GF150 buffer. The final concentrations in this mixture were TMR-dynein at 0.1 μM, dynactin at 0.2 μM, Lis1 at 6 μM and the adaptor (CDR2L1-159, CDR2L1-290, CDR2L1-290 ΔCC1 or JIP31-185) at 2 μM. Complexes were incubated on ice for 15 min then diluted with TIRF-dilution buffer (TIRF buffer supplemented with 75 mM KCl and 1 mg/mL casein) to a final volume of 10 μL. 1 μL of this complex was added to a mixture of 15 μL of TIRF-Casein buffer supplemented with 1 μL each of an oxygen scavenging system (4 mg/mL catalase, Merck; and 30 mg/mL glucose oxidase, Merck, dissolved in TIRF buffer), 45% (w/v) glucose, 30% BME, and 100 mM Mg-ATP. The final composition of this mixture was 25 mM HEPES pH 7.2, 4 mM MgSO_4_, 0.8 mM EGTA, 1.7 mM DTT, 45 mM KCl, 0.2 mg/mL catalase, 1.5 mg/mL glucose oxidase, 2.25% glucose, 1.5% BME, 5 mM ATP, 3.75 μM paclitaxel, 3 nM TMR-dynein, 6 nM dynactin, 60 nM adaptor and 180 nM LIS1. This mixture was flowed into the chamber. The sample was imaged immediately at 23°C using a TIRF microscope (Nikon Eclipse Ti inverted microscope equipped with a Nikon 100× TIRF oil immersion objective). For each sample, a microtubule image was acquired using a 488 nm laser. Following this a 500-frame movie was acquired (200 ms exposure, 4.1 fps) using a 561 nm laser. To analyse the data, ImageJ was used to generate kymographs from tiff movie stacks. Events of similar length were picked to analyse number of processive events/μm microtubule/min, using criteria outlined previously (Schlager *et al*., 2014; Urnavicius *et al*., 2018). Three or four technical replicates were performed for each sample.

### Cell culture

HeLa and HEK-293T cells were maintained at 37°C and 5% CO_2_ in high glucose Dulbecco’s modified Eagle’s medium (DMEM) supplemented with 10% fetal bovine serum, GlutaMAX, and 1% penicillin/streptomycin (all reagents from Gibco). Cell lines were regularly tested for mycoplasma contamination by PCR.

### Lentivirus production

HEK-293T cells were seeded 24 hour prior to transfection in 6-well plates at a density of 7×10^5^ cells/mL. Lentivirus was produced by co-transfecting cells with 1.2 µg transfer plasmid (pLenti-sgRNA or pKM808 containing protospacer sequences targeting CDR2 or CDR2L; pLenti-CMV-GFP-Hygro containing 3xFLAG::CDR2 or 3xFLAG::CDR2L), 0.3 µg envelope plasmid pMD2.G (Addgene 12259) and 1 µg packaging plasmid psPAX2 (Addgene 12260) using Lipofectamine 2000. The medium was changed 24 hours after transfection. Culture supernatant containing the lentivirus was collected 72 hours after transfection and stored for 24 hours at -80°C before transduction.

### CRISPR/Cas9-mediated genome editing and transgenic cell lines

To generate CDR2/CDR2L single and double KO cells, a HeLa cell line containing doxycycline-inducible human codon-optimized spCas9 was used (McKinley *et al*., 2015). For transduction with lentivirus, 400 μL virus-containing supernatant were added to 5×10^5^ cells suspended in 600 µL per well in a 24-well plate. Polybrene was added to a final concentration of 10 μg/mL, and the cell suspension was centrifuged in the 24-well plate at 1200 rcf (slow acceleration and deceleration) for 45 min at 37°C in a Mega Star 4.0R centrifuge with a TX-1000 rotor. Viruses were removed 24 hours later, and after a further 24 hours, antibiotics (1 µg/mL puromycin or 5 µg/mL blasticidin S) were added for 6–10 days. Cas9 expression was then induced with 1 µM doxycycline for 3 days. Colonies derived from single cells were obtained by seeding ∼100 cells in a 10-cm dish and allowing colonies to grow for 15 days. Individual colonies were collected by small-scale trypsinization and clones were expanded and screened by immunoblotting with antibodies against CDR2 and CDR2L. Single KO cells were generated first, and CDR2/L double KO cells were subsequently generated by targeting CDR2 in CDR2L KO cells.

To generate cells stably expressing GFP::3xFLAG-tagged CDR2 or CDR2L, CDR2/L double KO cells were infected with corresponding lentivirus and selected with 400 µg/µL hygromycin B for 12 days. Clonal lines were obtained as described above.

### Transient expression

24 hours prior to transfection, cells were seeded in a 24-well plate at 60,000 cells/well. For each well, 250 ng plasmid DNA and 0.75 µL Lipofectamine 2000 (Invitrogen) were combined in total of 100 µL Opti-MEM (Gibco) and incubated for 20 min at room temperature. The DNA–lipid complexes were then added to the well in a dropwise manner. After 24 hours, cells were processed for immunofluorescence as described below.

### RNA interference

24 hours prior to transfection, cells were seeded in a 24-well plate at 20,000 cells/well in medium without antibiotics. Cells were transfected with siRNAs (Dharmacon On-TARGETplus; Horizon Discovery) targeting KTN1 (SMARTpool J-010605-05-08), eEF1Bβ (SMARTpool J-011648-05-08), or Luciferase GL2 Duplex (D-001100-01) as a control. For each transfection, 1 μL of Lipofectamine RNAi-MAX (Invitrogen) and 50 nM of each siRNA were diluted in a total of 100 μL Opti-MEM and incubated for 20 min at room temperature. The siRNA-lipid complexes were then added in a dropwise manner to cells. After incubation for 6 hours, the transfection mixture was replaced with fresh complete medium, and cells were processed for immunofluorescence or immunoblotting 72 hours later.

### Immunofluorescence

Cells grown on 13-mm round coverslips (No. 1.5H, Marienfeld) coated with poly-L-lysine were fixed with 4% paraformaldehyde (PFA), diluted from a 20% aqueous solution (Delta Microscopies), in PBS for 30 min at room temperature and permeabilized with 0.1% (v/v) Triton X-100 in PBS for 10 min. Autofluorescence was quenched with 20 mM glycine in PBS for 10 min, and cells were incubated with blocking solution (3% (w/v) BSA in PBS) for 30 min. Coverslips were placed in a humid chamber and cells were incubated overnight at 4°C with the following primary antibodies diluted in blocking solution: mouse monoclonal anti-FLAG clone M2 (Merck F1804; 1:1000), rabbit monoclonal anti-KTN1 clone D5F7J (Cell Signaling Technology #13243; 1:100), rabbit polyclonal anti-CDR2 (Merck HPA023870; 1:200), mouse monoclonal anti-Climp63 clone G1/296 (MyBioSource MBS567120; 1:500), mouse monoclonal anti-GFP clone 9F9.F9 (Abcam ab1218; 1:500), rabbit monoclonal anti-Calnexin clone C5C9 (Cell Signaling Technology #2679; 1:100), mouse monoclonal anti-EF-1δ clone A-5 (Santa Cruz Biotechnology sc-393731; 1:200), mouse monoclonal anti-CETN3 clone 3E6 (Abnova H00001070-M01; 1:500). Coverslips were washed 3 × 5 min with PBS and incubated for 1 hour at room temperature in blocking solution containing the following donkey polyclonal secondary antibodies conjugated to Alexa dyes (Jackson ImmunoResearch; 1:300): anti-mouse IgG Alexa 488 (715-545-150), anti-mouse IgG Alexa 594 (715-585-150), anti-rabbit IgG Alexa 488 (711-545-152) and anti-rabbit IgG Alexa 594 (711-585-152). Coverslips were washed 3 × 5 min in PBS, rinsed once in H_2_O and mounted in ProLong Gold Antifade Mountant with DAPI (Thermo Fisher Scientific).

Cells were imaged on an Axio Observer microscope (Zeiss) equipped with an Orca Flash 4.0 camera (Hamamatsu) and an HXP 200C Illuminator (Zeiss), controlled by ZEN 2.3 software (Zeiss). Image stacks were acquired with a step size of 0.24 µm using a 63× NA 1.4 Plan-Apochromat objective. For presentation, images were pseudo-colored, cropped, and linearly adjusted for contrast using Fiji software (ImageJ2, version 2.14.0/1.54f). Images shown in figures correspond to individual images from a z-stack, unless stated otherwise. For quantification (Fig. 3A, D, G; Fig. 5C; Fig. S3C), images were acquired randomly using identical settings for the different conditions in an experiment, and maximum intensity projections of z-stacks were used to score ER morphology. Cells were classified as having clustered ER if the signal was densely concentrated adjacent to the nucleus, extending along less than half of the nuclear circumference. Cells were classified as containing ER patches when they contained one or more irregularly shaped areas of at least 3 µm^2^ with bright signal. Most CDR2/L double KO cells scored as "patchy" were well above this threshold, typically containing multiple patches of up to 15 µm^2^ in size.

### Immunoblotting

Cells grown in 24-well plates were collected by scraping with a pipette tip in 60 µL 1× SDS-PAGE sample buffer. Samples were heated for 3 min 95°C, vortexed, and centrifuged in an Eppendorf 5424 at 20,000 rcf for 5 min at room temperature. Proteins were resolved by 10 or 12% SDS-PAGE and transferred to a 0.2-µm nitrocellulose membrane (GE Healthcare). The membrane was blocked with 5% non-fat dry milk in TBS-T (20 mM Tris-HCl pH 7.5, 140 mM NaCl, 0.2% (v/v) Tween 20) for 1 hour and probed overnight at 4°C with the following primary antibodies diluted in 5% non-fat dry milk/TBS-T: mouse monoclonal anti-p150 clone 1/p150Glued (BD Transduction Laboratories 610473; 1:2500), mouse monoclonal anti-DIC clone 74.1 (Dillman and Pfister, 1994; 1:5000), mouse monoclonal anti-GAPDH clone 1E6D9 (Proteintech 60004-1-Ig; 1:5000), mouse monoclonal anti-α-tubulin clone B-5-1-2 (Merck T5168; 1:5000), mouse monoclonal anti-EF-18 clone A-5 (Santa Cruz Biotechnology sc-393731; 1:2000), rabbit monoclonal anti-KTN1 clone D5F7J (Cell Signaling Technology #13243; 1:2000), rabbit polyclonal anti-CDR2 (Merck HPA023870; 1:1000), rabbit polyclonal anti-CDR2L (Proteintech 14563-1-AP; 1:2000). The membrane was washed 4 × 7 min with TBS-T and incubated for 1 hour at room temperature in 5% non-fat dry milk/TBS-T containing goat polyclonal anti-mouse IgG (115-035-003) or anti-rabbit IgG (111-035-003) coupled to horseradish peroxidase (Jackson ImmunoResearch; 1:10000). The membrane was washed again 4 × 7 min with TBS-T and incubated with Pierce ECL Western Blotting Substrate (Thermo Fisher Scientific 32106) or Clarity Western ECL Substrate (for CDR2 and CDR2L antibodies; Bio-Rad 1705061). Proteins were visualized using Amersham Hyperfilm ECL (Cytiva) or the Bio-Rad ChemiDoc XRS+ system controlled by Image Lab software.

The intensity of protein bands in images acquired by the ChemiDoc XRS+ system (Fig. 5A) were quantified using Fiji software. The final integrated intensity of the band was calculated by subtracting the integrated intensity of a background region of the same size adjacent to the band. For each immunoblot, the eEF1Bβ signal was normalized to the α-tubulin signal, while KTN1 and CDR2 signals were normalized to the GAPDH signal. The normalized signal in the Luciferase RNAi condition was set to 1 in each experiment.

### Immunoprecipitation and mass spectrometry

CDR2/L double KO cells (control) and CDR2/L double KO cells expressing GFP::3xFLAG::CDR2 or GFP::3xFLAG::CDR2L were grown to 90% confluency in 40 15-cm dishes. To each dish, 4 mL PBS with 3 mM EDTA were added for 5 min at room temperature, and cells were harvested with a cell scraper and collected into 50-mL tubes. Cells were pelleted at 185 rcf with slow deceleration for 5 min at 4°C in a Mega Star 4.0R centrifuge with a TX-1000 rotor, washed sequentially with 50 mL PBS and 10 mL freezing buffer (50 mM HEPES pH 7.5, 100 mM KCl, 1 mM MgCl_2_, 1 mM EGTA, 10% (v/v) glycerol and 0.05% (v/v) NP-40), resuspended in 1 mL freezing buffer, flash-frozen in liquid nitrogen in a dropwise manner, and stored at -80°C.

Two replicate immunoprecipitations were performed per condition (on separate days). For each immunoprecipitation, half of the frozen cell droplets were thawed with 3 mL lysis buffer (freezing buffer supplemented with cOmplete EDTA-free protease inhibitor cocktail (1 tablet per 10 mL), 5 mM β-glycerophosphate, and 200 nM microcystin) in a 5-mL tube and lysed by sonication using a Branson sonifier 250 with a micro tip. The lysate was split equally into two 2-mL tubes and cleared at 20,000 rcf for 10 min at 4°C in a Megafuge 8R. The cleared lysate was transferred to new 2-mL tubes containing 70 µL Anti-FLAG M2 affinity gel (Merck) pre-eluted with

0.1 M glycine pH 2.6 and equilibrated with lysis buffer. The resin/lysate mixture was rotated for 1 hour at 4°C, transferred to a gravity flow column, and the resin was washed with 3 × 1 mL ice-cold lysis buffer containing 300 KCl and with 2 × 1 mL lysis buffer/300 KCl without NP-40. Proteins were eluted with 3 × 150 µL 0.1 M glycine pH 2.6 into a 1.5-mL tube containing 150 µL 2 M Tris-Cl pH 8.5. Proteins were precipitated with 20% trichloroacetic acid overnight on ice.

For LC-MS, proteins were reduced, alkylated and digested with trypsin following the solid-phase-enhanced (SP3) sample preparation approach (Hughes *et al*., 2019). Data was acquired on an Ultimate 3000 liquid chromatography system connected to a Q-Exactive mass spectrometer (Thermo Scientific), as described in Osório *et al*. (2021). Proteins were identified with Proteome Discoverer software v3.0.1.27 (Thermo Scientific) using the UniProt database (*Homo sapiens* proteome, 20,389 entries, 2022_05). Relative protein abundances between samples were determined using the label-free quantification (LFQ) method. Only proteins with a minimum of two unique peptides or two razor peptides and an abundance count of at least 10 were considered.

### Transmission electron microscopy

For TEM analysis, cells grown on 13-mm round poly-L-lysine-coated coverslips were fixed by adding to the culture medium an equal volume of 4% PFA (Electron Microcopy Sciences) and 5% glutaraldehyde (GTA) (Electron Microscopy Sciences) in 0.2 M cacodylate pH 7.4 for 15 min at room temperature. The fixation medium was removed, cells were further fixed with 2% PFA and 2.5% GTA in 0.1 M cacodylate pH 7.4 for 1 hour and washed 3 times with 0.1 M cacodylate pH 7.4. Cells were incubated with 1% osmium tetroxide (Electron Microscopy Sciences) in 0.1 M cacodylate pH 7.4 for 1 hour, washed 3 times with H_2_O, incubated with 1% uranyl acetate (Electron Microscopy Sciences) for 30 min, washed again with H_2_O, dehydrated through graded series of ethanol (50-70-80-100-100-100%), and embedded in Embed-812 resin (Electron Microscopy Sciences).

Ultrathin sections (70 nm) were cut on an RMC Ultramicrotome (PowerTome) using a diamond knife and recovered to 200 mesh nickel grids (Electron Microscopy Sciences), followed by post-staining with UranylLess (Electron Microscopy Sciences) and 3% lead citrate solution (Electron Microscopy Sciences) for 5 min each. Images were acquired at 80 kV on a JEM 1400 transmission electron microscope (JEOL) equipped with a PHURONA CMOS camera (EMSIS). For each condition (Fig. 3B; Fig. S3D), images were taken randomly in a section approximately corresponding to the central plane of cell nuclei.

Images captured at 3000–6000× magnification, enabling visualization of entire cells, were used for the quantification of ER sheet stacks (Fig. 3B; Fig. S3D). For each cell, the stack containing the greatest number of sheets was identified, and the number of stacked sheets was documented. Each condition was repeated three times with 40–50 cells scored per replicate.

### Correlative light–electron microscopy

Cells were grown in 35-mm glass bottom dishes (P35G-1.5-14-C-Grid, MatTeK) coated with poly-L-lysine. Cells were fixed with PFA and GTA as described for TEM, washed with PBS, permeabilized with 0.1% (w/v) saponin in PBS for 10 min, and incubated in PBS/0.1% saponin containing 20 mM glycine for 10 min. Cells were then incubated in blocking solution (see immunofluorescence) supplemented with 0.1% saponin for 30 min at room temperature. Anti-KTN1 antibody (see immunofluorescence) was diluted in the same solution and added to cells overnight at 4°C in a humid chamber. Cells were washed 3 × 5 min with PBS/0.1% saponin, and incubated with Alexa 594-conjugated secondary antibody (Jackson ImmunoResearch 711-585-152) diluted in blocking buffer. Cells were washed 3 × 5 min with PBS and imaged in PBS on a Nikon Eclipse Ti microscope coupled to an Andor Revolution XD spinning disk confocal system, composed of an iXon Ultra 897 CCD camera (Andor Technology), a solid-state laser combiner (ALC-UVP 350i, Andor Technology), and a CSU-X1 confocal scanner (Yokogawa Electric Corporation), controlled by Andor IQ3 software (Andor Technology). A z-stack (0.1 µm step size) through the entire cell was acquired with a 100x NA 1.45 Plan-Apochromat objective (Nikon). After fluorescence imaging, cells were further processed for TEM as described above, except that sequential sections were cut at 70 nm and formvar-coated slot grids (Electron Microscopy Sciences) were used. Fluorescence images corresponding to EM images were identified based on the shape of the cell’s outer boundary. Images were linearly resized, rotated, and moved in x and y to achieve best visual overlay using Fiji software.

### Structure prediction

The ColabFold implementation (Mirdita *et al*., 2022) of AlphaFold2 (Jumper *et al*., 2021) was used for structure prediction. The CDR2–DLIC1–DHC–DIC2 complex (Fig. S1A) was predicted by running ColabFold v1.5.5 with default parameters on two copies of CDR2 1–139 (UniProt Q01850), two copies of DC1L1 440–455 (UniProt Q9Y6G9), one copy of DYHC1 576–864 (UniProt Q14204) and one copy of DC1I2 226–583 (UniProt Q13409). The CDR2–KTN1 complex (Fig. 2D) was predicted using two copies each of CDR2 421–454 (UniProt Q01850) and KTN1 1114–1357 (UniProt Q86UP2). The eEF1Bβ–KTN1 complex (Fig. S3F) was predicted using one copy of EF1D 1–281 (UniProt P29692) and two copies of KTN1 1114–1357 (UniProt Q86UP2). The structure showing that eEF1Bβ and CDR2 occupy the same binding site on KTN1 (Fig. 4A) was predicted using one copy of CDR2 421–454 (UniProt Q01850), one copy of EF1D 39–68 (UniProt P29692) and two copies of KTN1 1114–1357 (UniProt Q86UP2). Structures were visualized with UCSF ChimeraX (Pettersen *et al*., 2021).

### Graphs and statistical analysis

Prism 10.0 software (GraphPad) was used for statistical analysis and to generate graphs. Statistical significance was determined using a two-tailed t test or ordinary one-way ANOVA followed by Tukey’s multiple comparisons test. The analytical method used is specified in the figure legends.

## ACKNOWLEDGEMENTS

This project was funded by the Fundação para a Ciência e a Tecnologia (FCT)/Ministério da Ciência, Tecnologia e Ensino Superior (PTDC/BIA-CEL/1321/2021). A.C., C.M.C.A., R.G. and T.J.D. were supported by FCT positions CEECIND/01967/2017, CEECIND/00771/2017, CEECIND/00333/2017 and CEECIND/01985/2018, respectively. V.T. was supported by FCT PhD fellowship SFRH/BD/147283/2019. Mass spectrometry was performed at the Proteomics i3S Scientific Platform with the assistance of Hugo Osório and support from the Portuguese Mass Spectrometry Network, integrated in the National Roadmap of Research Infrastructures of Strategic Relevance (ROTEIRO/0028/2013; LISBOA-01-0145-FEDER-022125). Electron microscopy was performed at the Histology and Electron Microscopy i3S Scientific Platform with the assistance of Rui Fernandes and Sofia Pacheco. UCSF ChimeraX is developed by the Resource for Biocomputing, Visualization, and Informatics at the University of California, San Francisco, with support from National Institutes of Health R01-GM129325 and the Office of Cyber Infrastructure and Computational Biology, National Institute of Allergy and Infectious Diseases.

The authors declare no competing financial interests.

---

**Figure S1:**
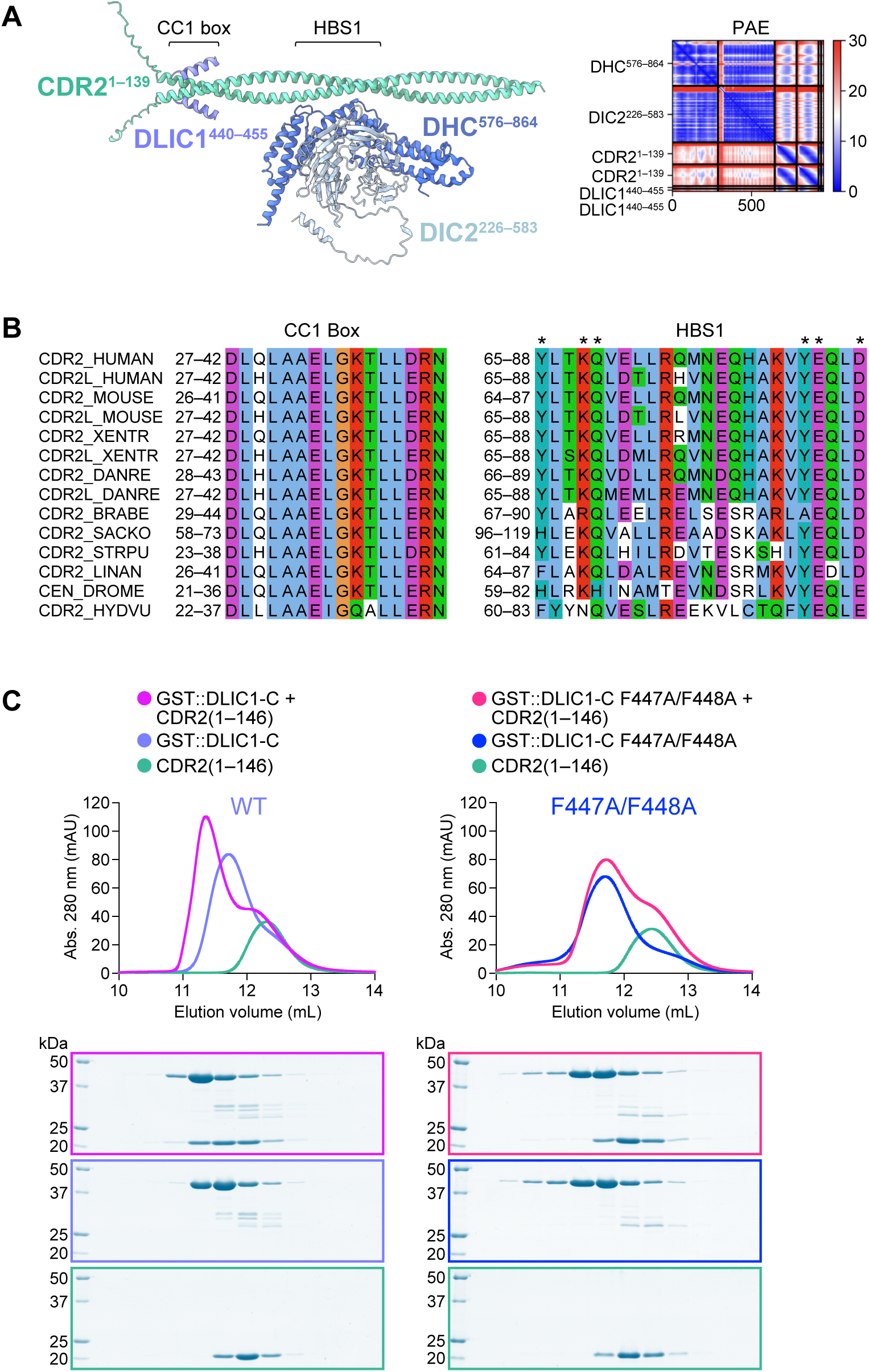
Related to Figure 1. **(A)** AF2 model and PAE plot of the CDR2 N-terminal coiled-coil in complex with the DLIC1 C-terminal helix and an N-terminal DHC fragment, which in turn is bound to the WD40 domain of DIC2. **(B)** Sequence alignment of the CC1 box and the dynein heavy chain binding site 1 (HBS1) in CDR2 and CDR2L proteins from different species (note invertebrates possess a single CDR2/CDR2L homolog). The HBS1 sequence is divergent from that of other adaptors but the interaction is predicted at the correct distance from the CC1 box. 6 residues, marked with asterisks, were mutated to alanine (HBS1_6A mutant) based on sequence conservation among CDR2 proteins and their position in the predicted structure. Accession numbers: CDR2_HUMAN (UniProt Q01850), CDR2L_HUMAN (UniProt Q86X02), CDR2_MOUSE (UniProt P97817), CDR2L_MOUSE (UniProt A2A6T1), CDR2_XENTR (UniProt F6R4S1), CDR2L_XENTR (UniProt A0A803JSM3), CDR2_DANRE (UniProt E7FC97), CDR2L_DANRE (UniProt Q6NZT2), CDR2_BRABE (UniProt A0A6P4ZS94), CDR2_SACKO (NCBI Reference Sequence XP_002736317.2), CDR2_STRPU (UniProt A0A7M7NRE1), CDR2_LINAN (NCBI Reference Sequence XP_013392376.1), CEN_DROME (UniProt Q9VIK6), CDR2_HYDVU (UniProt A0A8B6XII3). Species key (Phylum): HUMAN, *Homo sapiens* (Chordata); MOUSE, *Mus musculus* (Chordata); XENTR, *Xenopus tropicalis* (Chordata); DANRE, *Danio rerio* (Chordata); BRABE, *Branchiostoma belcheri* (Chordata); SACKO, *Saccoglossus kowalevskii* (Hemichordata); STRPU, *Strongylocentrotus purpuratus* (Echinodermata); LINAN, *Lingula anatina* (Brachiopoda); DROME, *Drosophila melanogaster* (Arthropoda); HYDVU, *Hydra vulgaris* (Cnidaria). **(C)** Elution profiles and BlueSafe-stained SDS-PAGE gels of purified recombinant human CDR2 and DLIC1 fragments after SEC. DLIC1-C corresponds to residues 388-523. The elution profile and gel for CDR2 are shown on both left and right to facilitate comparison between wild-type DLIC1-C and the F447A/F448A mutant. Molecular weight is indicated in kilodaltons (kDa).

**Figure S2:**
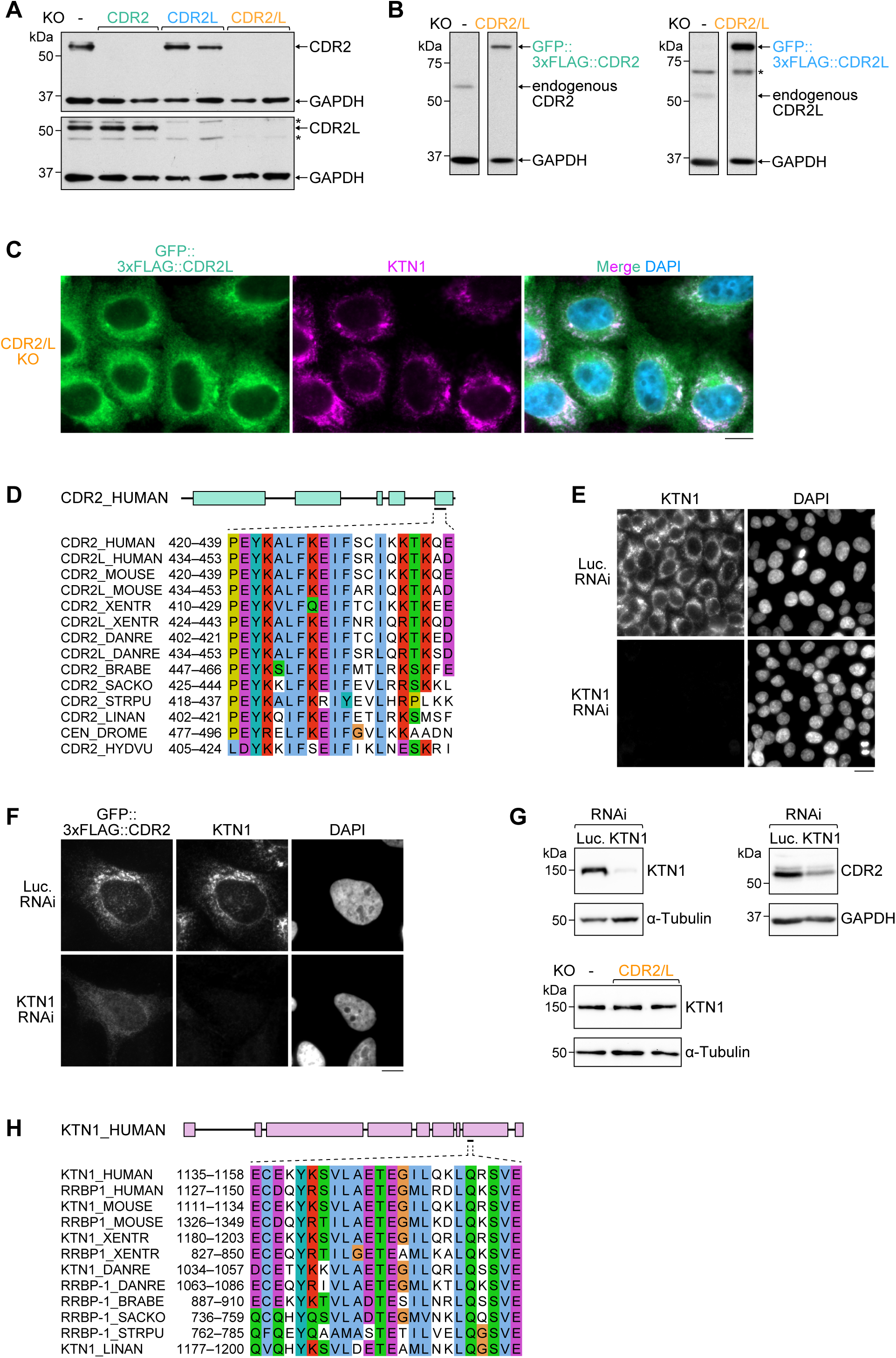
Related to Figure 2. **(A)** Immunoblots of HeLa cells harboring single and double KOs of CDR2 and CDR2L (two independently derived cell lines were analyzed for each condition). GAPDH serves as the loading control. Molecular weight is indicated in kilodaltons (kDa). **(B)** Immunoblots of CDR2/L double KO cells stably expressing GFP::3xFLAG::CDR2 or CDR2L, used for the experiments in Fig. 2A. GAPDH serves as the loading control. Molecular weight is indicated in kilodaltons (kDa). **(C)** Immunofluorescence of CDR2/L double KO cells stably expressing GFP::3xFLAG::CDR2L, showing co-localization with KTN1 and diffuse cytoplasmic signal. Note that while average expression levels of transgene-encoded CDR2L are significantly higher than those of endogenous CDR2L, as shown in *(B)*, expression in individual cells is variable. Cells shown here have relatively low expression levels. Scale bar, 10 µm. **(D)** Sequence alignment of the C-terminal helix in CDR2 and CDR2L proteins from different species. Accession numbers and species key as in *Fig. S1B*. **(E) –(G)** Immunofluorescence images and immunoblots showing knockdown of KTN1 by RNAi and the resulting delocalization/destabilization of CDR2 in HeLa cells. By contrast, KTN1 levels remain unaffected in CDR2/L double KO cells (two independently derived KO cell lines were analyzed). Scale bars, 20 µm *(E)* and 10 µm *(F)*. Molecular weight is indicated in kilodaltons (kDa). **(H)** Sequence alignment of the CDR2/eEF1Bβ binding site in KTN1 and its paralog RRBP1 (p180) from different species (invertebrates possess a single KTN1/RRBP1 homolog). Accession numbers: KTN1_HUMAN (UniProt Q86UP2), RRBP1_HUMAN (Q9P2E9), KTN1_MOUSE (UniProt Q61595), RRBP1_MOUSE (UniProt Q99PL5), KTN1_XENTR (UniProt B3DL66), RRBP1_XENTR (UniProt F7A6K6), KTN1_DANRE (UniProt E7F049), RRBP1_DANRE (UniProt B8A4D7), RRBP1_BRABE (UniProt A0A6P5A3T7), RRBP1_SACKO (NCBI Reference Sequence XP_002741373.1), RRBP1_STRPU (A0A7M7LVI4), KTN1_LINAN (NCBI Reference Sequence XP_013397491.1). Species key as in *Fig. S1B*. No CDR2 binding site could be identified for the KTN1/RRBP1 homologs of DROME and HYDVU (UniProt Q960Y8 and T2M451, respectively), despite the presence of a well conserved CDR2 helix, as shown in *(D)*.

**Figure S3:**
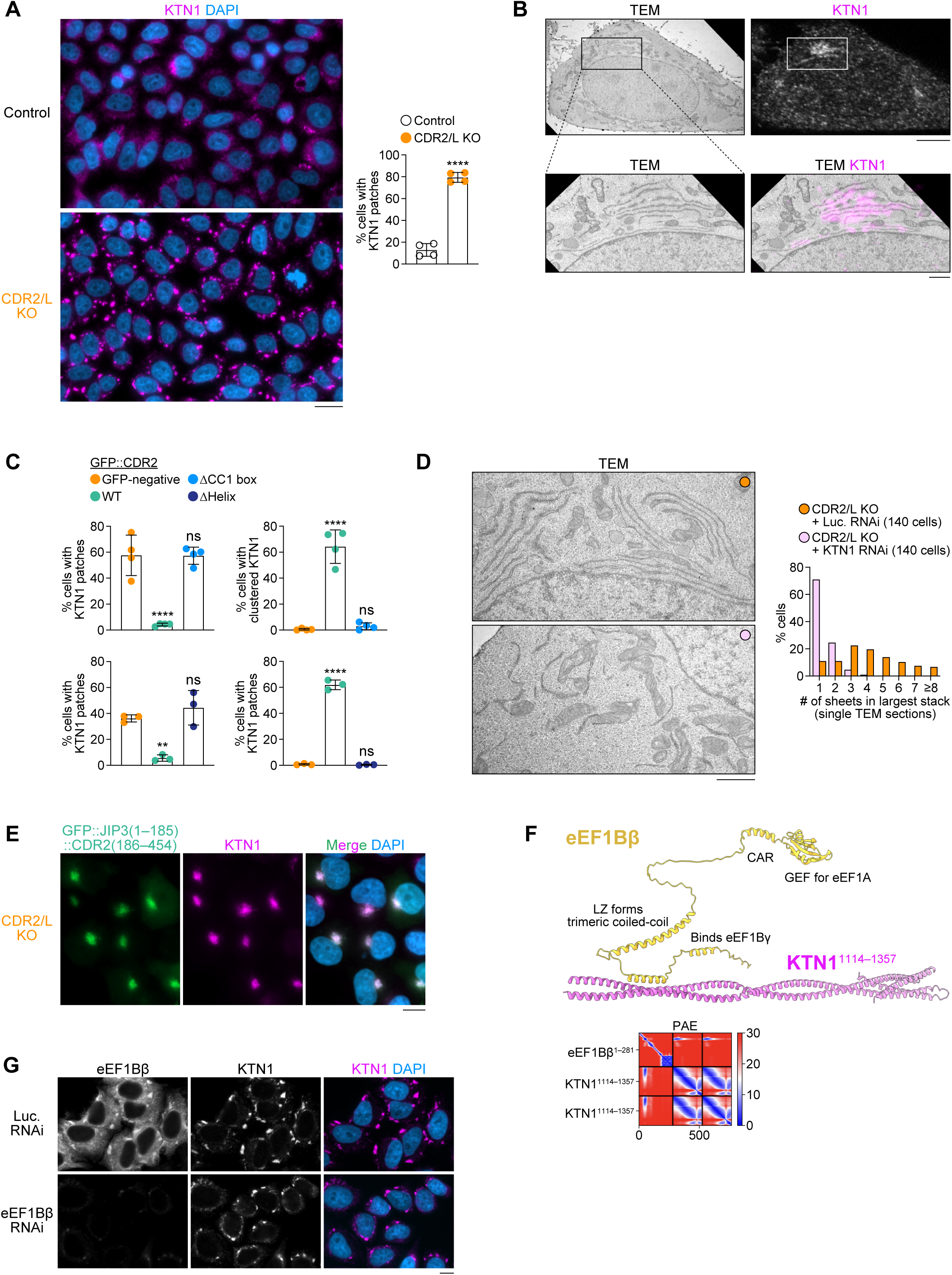
Related to Figures 3, 4 and 5. **(A)** *(left)* Immunofluorescence images showing exacerbated patchy distribution of KTN1 in HeLa CDR2/L double KO cells. Scale bar, 20 µm. *(right)* Fraction of cells with prominent KTN1 patches, plotted as mean ± SD (4 independent experiments, >1000 cells scored in total per condition). Statistical significance was determined using a two-tailed t test. \*\*\*\**P* < 0.0001. These cells were treated with siRNA against Luciferase, which further enhances KTN1 patch formation in CDR2/L double KO cells relative to untreated cells (compare with quantification in Fig. 3A). **(B)** Correlative light–electron microscopy images of CDR2/L double KO cells showing that the KTN1 patches observed by immunofluorescence correspond to stacked ER sheets. Scale bars, 5 µm *(top)* and 1 µm *(bottom)*. **(C)** Fraction of cells (mean ± SD, 4 and 3 independent experiments for ΔCC1 box and ΔHelix, respectively; >580 cell scored in total per condition) with prominent KTN1 patches *(left)* or centrosome-proximal KTN1 clustering *(right)* in the conditions shown in Fig. 3E, using a second independently derived CDR2/L double KO cell line. ΔCC1 box and ΔHelix experiments each have their own WT and GFP-negative controls. Statistical significance was determined using ordinary one-way ANOVA followed by Tukey’s multiple comparisons test. \*\*\*\**P* < 0.0001; \*\**P* < 0.01; *ns* = not significant, *P* > 0.05. **(D)** *(left)* TEM images of ER sheets in CDR2/L double KO cells with and without knockdown of KTN1. Scale bar, 1 µm. *(right)* Number of ER sheets in the largest stack per cell, determined as described in Fig. 3B. The CDR2/L double KO data is the same as in Fig. 3B. **(E)** Immunofluorescence image showing penetrant and tight clustering of KTN1 in the presence of JIP3(1–185)::CDR2(186–454). Scale bar, 10 µm. **(F)** AF2 model and predicted alignment error (PAE) plot of full-length eEF1Bβ in complex with the KTN1 C-terminus. One copy of eEF1Bβ was used for the prediction, but note that eEF1Bβ can form a trimer through its leucine zipper (LZ) domain (Bondarchuk *et al*., 2022). **(G)** Immunofluorescence images (maximum intensity projection of z-stack) showing that eEF1Bβ knockdown in CDR2/L double KO cells does not alter KTN1 distribution (see corresponding quantification in Fig. 5C). Scale bar, 10 µm.

